# Cell-to-cell heterogeneity in p38-mediated cross-inhibition of JNK causes stochastic cell death

**DOI:** 10.1101/151852

**Authors:** Haruko Miura, Yohei Kondo, Michiyuki Matsuda, Kazuhiro Aoki

## Abstract

The stress activated protein kinases (SAPKs), c-Jun N-terminal kinase (JNK) and p38, are important players in cell fate decisions in response to environmental stress signals. Crosstalk signaling between JNK and p38 is emerging as an important regulatory mechanism in the inflammatory and stress responses. However, it is still unknown how this crosstalk affects signaling dynamics, cell-to-cell variation, and cellular responses at the single-cell level. To address these questions, we established a multiplexed live-cell imaging system based on kinase translocation reporters to simultaneously monitor JNK and p38 activities with high specificity and sensitivity at single-cell resolution. Various stresses, such as anisomycin, osmotic stress, and UV irradiation, and pro-inflammatory cytokines activated JNK and p38 with various dynamics. In all cases, however, p38 suppressed JNK activity in a cross-inhibitory manner. We demonstrate that p38 antagonizes JNK through both transcriptional and post-translational mechanisms. This cross-inhibition of JNK appears to generate cellular heterogeneity in JNK activity after stress exposure. Our data indicate that this heterogeneity in JNK activity plays a role in fractional killing in response to UV stress. Our highly sensitive multiplexed imaging system enables detailed investigation into the p38-JNK interplay in single cells.

**One Sentence Summary:** Cross-inhibition by p38 generates cell-to-cell variability in JNK activity.

## Introduction

Environmental stresses and inflammatory signals lead to adaptive cellular responses of reversible cell cycle arrest or irreversible cell senescence and death. In general, modest damage causes cell cycle arrest or senescence, while overwhelming damage triggers apoptosis (*1, 2*). The stress inputs are sensed and transmitted to a number of intracellular signaling cascades such as the stress activating protein kinase (SAPK) pathway and p53 pathway, followed by induction of cyclin dependent kinase inhibitors and the caspase activating cascade (*3*). Several mechanisms have been proposed to explain the alternative cell fate decisions in a manner dependent on the signaling threshold (*4, 5*) and signaling dynamics (*6–9*).

Cell death is an irreversible and switch-like process to eliminate damaged cells. Single-cell studies with live cell imaging have revealed cell-to-cell variability in the timing and probability of cell death (*10*). Such fractional (incomplete) cell killing is a major obstacle of cancer treatment with chemotherapy and antibacterial treatment with antibiotics. In some circumstances, a subset of tumor cells harboring specific mutations survives treatment with chemotherapy, leading to the fractional killing (*11*). Meanwhile, intrinsic noise and stochastic fluctuation of gene expression cause an increase in cell-to-cell heterogeneity that is known as non-genetic heterogeneity, and eventually lead to resistance or persistence to chemotherapy for cancer cells or antibiotics for bacteria at the population level (*12–14*). More recently, fractional killing has been reported to arise from the cell-to-cell variance of intracellular signaling such as Caspase 8 activity (*15*) and p53 dynamics (*16*). However, it is unclear whether the fractional killing arises from a more upstream signaling pathway, namely, the stress-activating protein kinase (SAPK) pathway.

The SAPKs, c-Jun N-terminal kinase (JNK) and p38, play important roles in the cellular response to a wide range of environmental stresses and inflammatory signals (*17–19*). They are members of the mitogen-activated protein kinase (MAPK) family, which are activated in a three-tiered signaling cascade of sequentially activated protein kinases: MAPK kinase kinase (MAPKKK or MKKK), MAPK kinase (MAPKK or MKK), and MAPK. There are two MAPKKs that activate JNK, namely MKK4 and MKK7, while p38 is mainly activated by MKK3 and MKK6. In contrast to the limited number of MAPKKs, at least 15 MAPKKKs function upstream of p38 and/or JNK (*20*), allowing responses to diverse stress signals such as osmotic shock, oxidative stress, ultraviolet (UV) irradiation, heat shock, DNA damage, protein synthesis inhibitors, and pro-inflammatory cytokines. After activation, SAPKs phosphorylate their target substrates, including protein kinases, regulatory proteins, and various transcription factors, in order to exert their diverse functions (*20, 21*).

In this study, we present a multiplexed imaging system based on kinase translocation reporters (KTRs) (*22*), which allow us to monitor p38 and JNK activities with high specificity and sensitivity in single living cells. We employed this imaging system to record p38 and JNK activity dynamics in response to various stress stimuli at single cell resolution, and quantitated the cross-inhibition of JNK by p38 through post-translational and/or transcriptional mechanisms. We demonstrated that the cross-inhibition generates the cell-to-cell variation of JNK activity, thereby leading to fractional killing upon UV irradiation.

## Results

### Development of a multiplexed imaging system for p38 and JNK activities

For the co-visualization of JNK and p38 kinase activities, we sought to implement a multiplexed imaging system based on genetically encoded KTRs. This approach depends on conversion of the phosphorylation event into a nucleocytoplasmic shuttling event. Such nucleocytoplasmic translocations have been described in some naturally occurring proteins (*23–26*), and more recently in the KTR technology (*22*). Upon phosphorylation, the reporter is exported from the nucleus to the cytosol, and goes back to the nucleus upon dephosphorylation (Fig. 1A). The kinase activity can be quantified by the ratio of the cytosolic to the nuclear fluorescence intensity (C/N ratio).

**Fig. 1.**
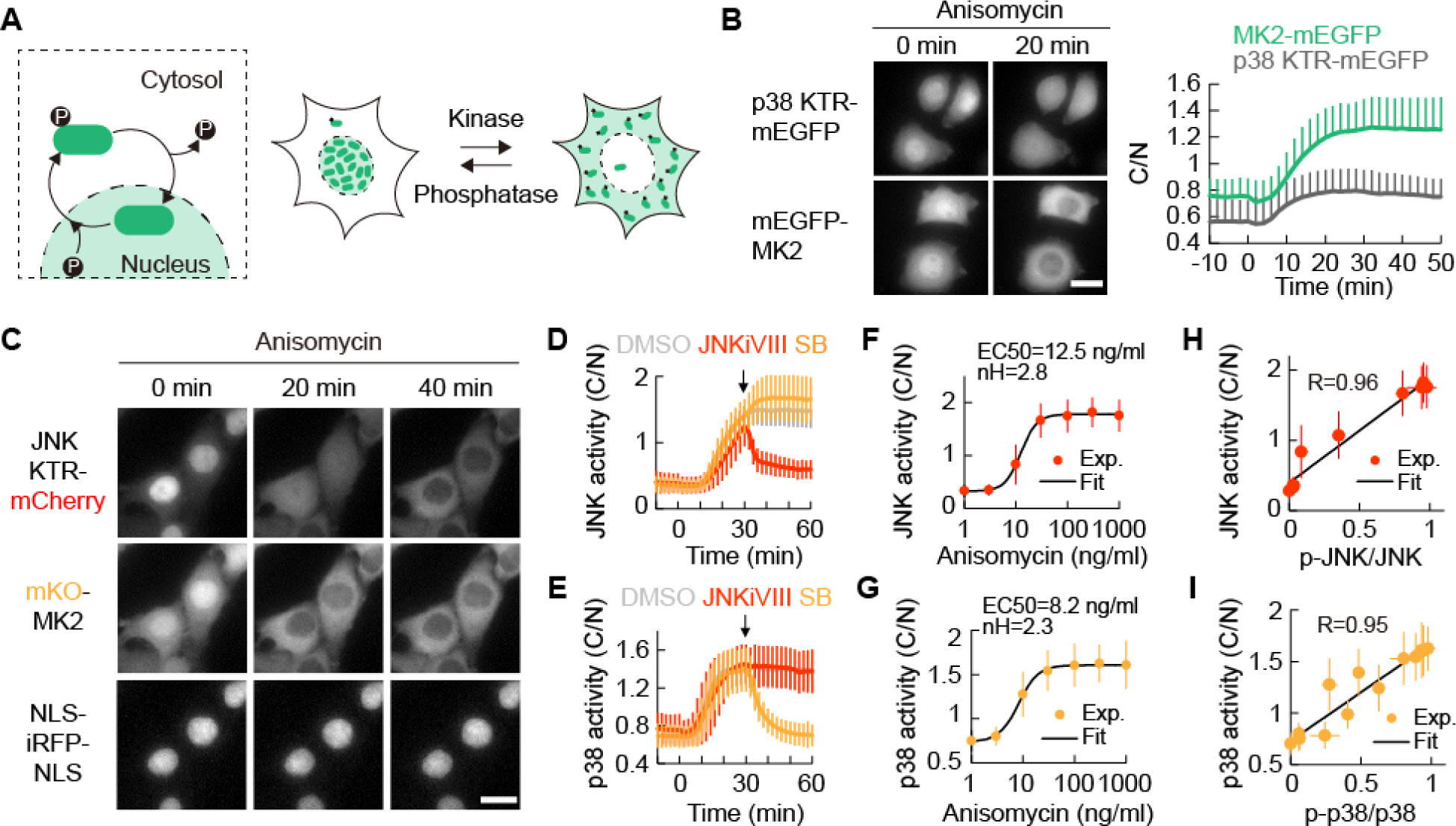
Establishment of a multiplexed imaging system for monitoring JNK and p38 kinase activities. (**A**) Schematic of the phosphorylation-mediated translocation of a kinase translocation reporter (KTR). (**B**) HeLa cells stably expressing p38 KTR-mEGFP and mEGFP-MK2, respectively, were stimulated with 1 μg/ml anisomycin. Representative images before and after 20 min of anisomycin stimulation and averaged C/N ratios with SD are shown (left). Scale bar: 20 μm. C/N ratios are plotted as a function of time (right). n = 38 cells for p38 KTR-mEGFP, n = 34 cells for mEGFP-MK2 from two independent experiments. (**C**) Clonal HeLa cells stably expressing JNK KTR-mCherry, mKO-MK2, and NLS-iRFP-NLS were stimulated with 1 μg/ml anisomycin and imaged over time. Scale bar: 20 μm. (**D** and **E**) Clonal HeLa cells stably expressing JNK KTR-mCherry, mKO-MK2, and NLS-iRFP-NLS were stimulated with 1 μg/ml anisomycin and treated with 10 μM JNK inhibitor VIII (JNKiVIII), 10 μM of the p38 inhibitor SB203580 (SB), or 0.1% DMSO at the indicated time point (arrow). Averaged JNK KTR-mCherry (D) and mKO-MK2 (E) C/N ratios with SD are shown. n = 40 cells for each condition from two independent experiments. (**F** and **G**) Dose-response curves of the JNK KTR-mCherry (F) and mKO-MK2 (G) C/N ratios after 1 h of anisomycin stimulation are shown. Experimental data were fitted with the Hill function and the derived EC50 values and Hill coefficients (nH) are indicated. Data are presented as the mean with SD. n ≥ 100 cells for each condition, from at least three independent experiments. (**H** and **I**) JNK KTR-mCherry (H) and mKO-MK2 (I) C/N ratios linearly correlate with normalized p-JNK/JNK and p-p38/p38 ratios, respectively. Imaging and western blot data were obtained after 1 h of treatment with various doses of anisomycin and/or SB203580. Scatterplots were fitted to a linear regression and the obtained Pearson correlation values are shown. Data are presented as the mean with SD. n = 3 cells for western blot data and n ≥ 60 cells for imaging data.

In a preliminary experiment, we used the JNK and p38 KTRs reported previously (*22*). However, in our hands the prototype p38 KTR showed limited cytosolic translocation by anisomycin stress, which is a strong inducer of the SAPK pathways (Fig. 1B), prompting us to develop an improved translocation-based reporter for p38 activity. It has been reported that the p38α substrate MAPKAPK2 (MK2) naturally shuttles between the nucleus and cytosol based on its phosphorylation status (*27–29*). We found that mEGFP-MK2, a full-length MK2 fused to monomeric EGFP (mEGFP), outperformed p38 KTR in terms of the increased C/N ratio in response to anisomycin (Fig. 1B and fig. S1A). Thus, hereafter, we used the fluorescent protein-tagged MK2 as the quantitative live cell imaging reporter for p38α activity.

For multiplexed imaging of the JNK and p38 activities, we fused JNK KTR to mCherry and MK2 to monomeric Kusabira Orange (mKO), respectively. In addition, a near-infrared fluorescent protein (iRFP) flanked by nuclear localization signals (NLS) was co-expressed as a nuclear marker. To examine the level of cross-excitation and/or bleed-through during the imaging of JNK and p38 reporters, we imaged HeLa cells expressing each reporter with all combinations of excitation/emission filters (fig. S1B). The bleed-through and cross-excitation of JNK KTR-mCherry, mKO-MK2, and NLS-iRFP-NLS were negligible under our imaging conditions, allowing multiplexed imaging of these reporters without linear unmixing.

To minimize the effect of reporter concentrations on the output, we established HeLa cell lines stably-expressing all three reporters by single cell cloning. We did not find a remarkable difference among the established cell lines and chose one of the cell lines that exhibited the expected response to various stimuli for a thorough analysis. After stimulation with 1.0 μg/ml anisomycin, translocation of JNK KTR-mCherry and mKO-MK2 was readily observed (Fig. 1C) and was quantified as the C/N ratio (Fig. 1, D and E, grey lines). The mKO-MK2 and JNK KTR-mCherry C/N ratios started to increase approximately 5 and 10 min after stimulation, respectively. These C/N ratios reached the plateau about 20 and 30 min after stimulation, respectively (Fig. 1, D and E). The kinetics of the JNK KTR-mCherry and mKO-MK2 C/N ratios were comparable to those of JNK and p38 phosphorylation examined by immunoblotting (fig. S1C). Responses to inhibitors were generally to be expected; the anisomycin-stimulated increase in the C/N ratio of JNK KTR-mCherry was specifically repressed by JNK inhibitor VIII, but not by the p38 inhibitor SB203580 (Fig. 1D), whereas the increase in the C/N ratio of mKO-MK2 was rapidly reduced only by SB203580, but not by JNK inhibitor VIII (Fig. 1E).

Interestingly, however, p38 inhibition increased the C/N ratio of JNK KTR-mCherry (Fig. 1D, blue line), an effect whose detailed mechanism will be described later. The specificity of the probes was further confirmed by the gene knock-out (KO) experiment. The anisomycin-induced increase in the C/N ratio of JNK KTR-mCherry was fully suppressed in *MAPK8* (*JNK1*) and *MAPK9* (*JNK2*) double KO cells, while translocation of mKO-MK2 was abolished in *MAPK14* (*p38α*) KO cells (fig. S1, D and E).

To evaluate the sensitivity and dynamic range of JNK KTR-mCherry and mKO-MK2, we examined the dose response to anisomycin. The C/N ratios of mKO-MK2 and JNK KTR-mCherry showed modest switch-like responses (Fig. 1, F and G); the EC50 values (sensitivities) were 8.2 ng/ml and 12.5 ng/ml, and the Hill coefficients were 2.3 and 2.8 for mKO-MK2 and JNK KTR-mCherry, respectively. Plotting the C/N ratios against the fraction of phosphorylated target kinases revealed a strong linear correlation, indicating that the dynamic range of the translocation reporters covers the full range of JNK and p38 activities upon anisomycin stimulation (Fig. 1, H and I, and fig. S1F). Taken together, the results showed that this system allows us to quantitatively monitor stress-induced p38 and JNK activity-dynamics with high specificity and sensitivity at the single cell level.

To facilitate the equimolar expression of reporter proteins, we next constructed a polycistronic vector, named pNJP (Nuclear, JNK, and p38 reporter), which comprised NLS-iRFP-NLS, JNK KTR-mCherry, and mKO-MK2, connected by self-cleaving P2A peptide sequences (*30*) (fig. S1G). HeLa cells stably expressing NJP showed the expected localizations (fig. S1H) and successful separation of the reporters (fig. S1I).

Of note, when JNK KTR-mCherry and mKO-MK2 were transiently overexpressed in HeLa cells, the increase in the C/N ratio was negatively correlated with the expression levels (fig. S1J, left); meanwhile, in the cells stably-expressing the reporters, the C/N ratio was independent of the reporter expression levels (fig. S1J, right).

### Crosstalk between JNK and the p38-signaling pathways

Prior work has provided compelling evidence of the crosstalk between p38 and JNK activities at the population level. Chemical inhibition or KO of *p38α* has led to JNK hyper-activation in several contexts, suggesting that p38 negatively regulates JNK (*31–38*). Although this relationship has been well established in the cell population with limited time points, it is virtually unknown how this crosstalk affects the dynamics and cell-to-cell variability of SAPK signaling at the single cell level. We chose this question to demonstrate the merit of the imaging system for JNK and p38 activities. The reporter HeLa cells pretreated with DMSO or SB203580 were stimulated with pro-inflammatory cytokines, 10 ng/ml tumor necrosis factor α (TNFα), 10 ng/ml interleukin-1β (IL-1β), or stress inputs, 10 ng/ml anisomycin, 200 mM sorbitol, and 100 J/m^2^ UV-C (Fig. 2A and fig. S2A). At the single cell level, we found that the JNK activity varied among individual cells, while the p38 response appeared more homogenous for all stimulants (Fig. 2A and Movie S1). When cells were pretreated with SB203580, JNK activity was significantly increased under all inflammatory and stress conditions, and its cellular heterogeneity seemed to be reduced (Fig. 2A).

**Fig. 2.**
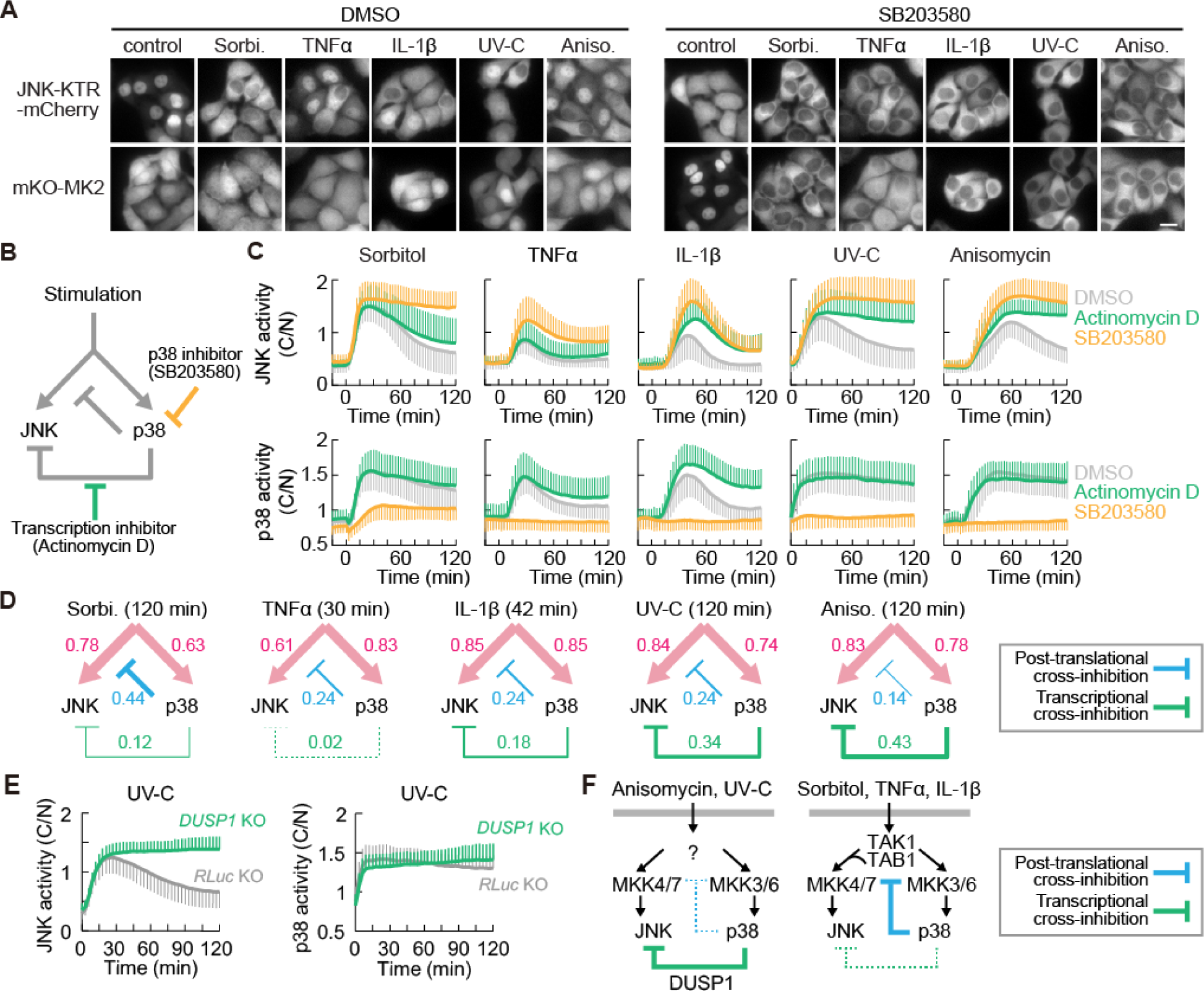
Transcriptional and post-translational cross-inhibition of JNK activity by p38 under various stress conditions. (**A**) Clonal HeLa cells stably expressing JNK KTR-mCherry, mKO-MK2, and NLS-iRFP-NLS were pretreated with 0.1% DMSO and 10 μM SB203580, respectively, and then stimulated with 10 ng/ml TNFα, 10 ng/ml IL-1β, 10 ng/ml anisomycin, 200 mM sorbitol, 100 J/m^2^ UV-C, or imaging medium as a control. Representative images of JNK KTR-mCherry and mKO-MK2 after 30 min of treatment with TNFα and 1 h of treatment with the other stimuli are shown. (**B**) Schematic representation of cross-inhibition mechanisms of JNK by p38. (**C**) Clonal HeLa reporter cells were pretreated with 0.1% DMSO, 1 μg/ml actinomycin D, or 10 μM SB203580, then stimulated with 200 mM sorbitol, 10 ng/ml TNFα, 10 ng/ml IL-1β, 100 J/m^2^ UV-C, or 10 ng/ml anisomycin, and imaged over time. Averaged JNK KTR-mCherry and mKO-MK2 C/N ratios with SD are shown. n = 70 cells per condition from at least two independent experiments. (**D**) JNK and p38 activation and post-translational and transcriptional cross-inhibition of JNK by p38 are indicated for 200 mM sorbitol at 120 min, 10 ng/ml TNFα at 30 min, 10 ng/ml IL-1β at 42 min, 100 J/m^2^ UV-C at 120 min, and 10 ng/ml anisomycin stimulation at 120 min. Edge thickness is proportional to the strength of pathway activities. (**E**) HeLa *DUSP1* KO and *RLuc* KO control cells stably expressing NJP were stimulated with 100 J/m^2^ UV-C. The time courses of the averaged JNK KTR-mCherry and mKO-MK2 C/N ratios with SD are shown. n = 50 cells for each condition from two independent experiments. (**F**) Schematic of post-translational and transcriptional cross-inhibition of JNK by p38. The TAK1/TAB1 complex might mediate cross-inhibition of JNK by p38, when stimulated with TNFα and IL-1β.

We next examined the mechanisms underlying the negative regulation of JNK by p38. Previous studies have demonstrated two possible mechanisms: Post-translational regulation of an upstream MAPKK/MAPKKK or transcriptional induction of phosphatases such as DUSP1. It is suggested that the mechanism underlying the cross-inhibition of JNK by p38 depends on the cell type and stimulus (*20, 39*). To dissect the post-translational and transcriptional mechanisms, we pretreated cells with an inhibitor of transcription, actinomycin D, or SB203580, followed by exposure to each stress input (Fig. 2B). SB203580 always enhanced stimulant-induced JNK activities more efficiently than actinomycin D (Fig. 2C, upper panels). Interestingly, actinomycin D markedly potentiated the JNK activation by IL-1β, anisomycin, or UV-C, but not the TNFα- and sorbitol-induced JNK activities (Fig. 2C, upper panels). Meanwhile, the transient p38 activation by cytokines (IL-1β and TNFα) was prolonged by actinomycin D, but the sustained p38 activation by the other stimulants was not affected (Fig. 2C, lower panels). The effect of SB203580 on JNK activity was confirmed by western blotting. Consistent with the imaging data, SB203580 pretreatment resulted in significantly higher JNK phosphorylation than the DMSO control under all the stress conditions examined (fig. S2B). SB203580 did not affect the phosphorylation of p38 itself (fig. S2B).

The heterogeneous sensitivity to SB203580 and actinomycin D enabled us to roughly quantify how much these two pathways contribute to the cross-inhibition of JNK by p38 under various stress and inflammatory conditions (Fig. 2D). We found that, in response to osmotic shock and TNFα, cross-inhibition is mainly mediated by post-translational mechanisms, while in the case of IL-1β, UV-C, and anisomycin stress, both modes of cross-inhibition seem to operate (Fig. 2D).

### Transcriptional and post-translational mechanisms of the cross-inhibition of JNK by p38

Earlier studies have demonstrated that p38 induces *DUSP1* expression, and thereby suppresses JNK (*33, 36, 40*). Indeed, we confirmed that *DUSP1* was increased within 45 min after the stimulation by anisomycin, IL-1β, and UV-C, and this increase was cancelled by actinomycin D (fig. S2C). The role played by DUSP1 was further examined in *DUSP1* KO cells (fig. S2D). Upon UV-C stimulation, *DUSP1* KO cells, but not control *RLuc* KO cells, showed sustained JNK activation. DUSP1 deficiency did not have any observable effect on p38 activation (Fig. 2E). Very similar results were obtained by the anisomycin treatment (fig. S2E). These data confirmed that DUSP1 is responsible for the p38-mediated transcriptional cross-inhibition of JNK, which plays a major role in cells stimulated with anisomycin and UV-C (Fig. 2F, left).

Next, we investigated the post-translational cross-inhibition mechanisms. TAB1 and TAK1 have been reported to be involved in this process (*31*) (Fig. 2F, right). As expected, we found that the TNFα- or IL-1β-induced JNK activities were almost completely suppressed in *TAB1* and *MAP3K7* (*TAK1*) KO cells (fig. S2, F and G). On the other hand, TNFα treatment showed that p38 activation was strongly dependent only on TAK1, and surprisingly, not on TAB1 (fig. S2G). IL-1β-induced p38 activation was partially suppressed in *TAK1* KO and *TAB1* KO cells (fig. S2G). The phosphorylation of TAB1 by p38 has been reported to be involved in JNK cross-inhibition upon TNFα and IL-1β stimulation (*31*). Therefore, we mutated the p38 phosphorylation sites in TAB1, Ser423 and Thr431, to alanine, and examined whether expression of this TAK1 SATA mutant rescued the cross-inhibition of JNK by p38 in *TAB1* KO HeLa cells. As expected, the TAB1 wildtype could rescue the cross-inhibition of JNK, whereas TAB1 SATA caused significantly higher JNK activities than did the TAB1 wildtype in response to TNFα and IL-1β (fig. S2G). In contrast, p38 activity seemed not to be affected by the SATA mutation (fig. S2G). These results support the previous observation that p38 inhibition or KO increased cytokine-induced TAK1 activity and JNK phosphorylation (*31*).

### Generation of heterogeneous JNK signaling among cells by p38-mediated cross-inhibition

So far, we have used population-based data to validate that the observations acquired by our reporter cells agree with previous biochemical results. To demonstrate the advantage of single cell analysis, we next asked how p38-mediated cross-inhibition of JNK generates cell-to-cell heterogeneity. As mentioned earlier, JNK activity varied more greatly among single cells after cytokine and stress exposure than did p38 activity (Fig. 2A and Movie S1). To quantitatively describe the cell-to-cell heterogeneity, we hereafter use the coefficient of variation (CV) of the distribution of C/N ratios. The distribution of activity 1 h after anisomycin stress (10 ng/ml) was broader for JNK, with a CV of ∼ 0.39, than for p38, with a CV of ∼ 0.21 (Fig. 3A). Remarkably, inhibition of p38 not only shifted the distribution of JNK activities to higher levels, but also reduced the variation of JNK activity to a CV of ∼ 0.18, which is approximately half the value of the control (Fig. 3A). This effect was also well represented in the time evolution of the CV upon stress stimulation; anisomycin treatment gradually increased the CV of JNK activity, while SB203580 reduced the CV of JNK activity after anisomycin stimulation (Fig. 3B). Consistently, *MAPK14* (*p38α)* KO reduced the cell-to-cell variability of JNK activity upon anisomycin stress (fig. S3, A and B). Of note, the CV values of JNK activity reached the minimum in SB203580-treated cells 30 min after anisomycin stimulation, even though the JNK activity had not yet reached its maximal level at that time (Fig. 2C and 3B). This result implies that the reduction of CV was not caused by the saturation of JNK activity. In the case of p38 activity, the CV remained constant over the time course of anisomycin stimulation (Fig. 3B). Similar results were obtained for other stimuli: SB203580 led to smaller CVs in JNK activity in response to 10 ng/ml TNFα, 10 ng/ml IL-1β, 200 mM sorbitol, and 100 J/m^2^ UV-C, in comparison to the control (Fig. 3C). In agreement with the results showing that p38-mediated *DUSP1* induction is a key module of cross-inhibition, *DUSP1* KO led to a clearly reduced CV of JNK activity in response to UV-C and anisomycin stress compared to the control (Fig. 3, D and E, and fig. S3, C and D). This observation points to the hypothesis that noise in the p38-mediated gene expression of *DUSP1* might be a potential source of the cell-to-cell variation of JNK activity.

**Fig. 3.**
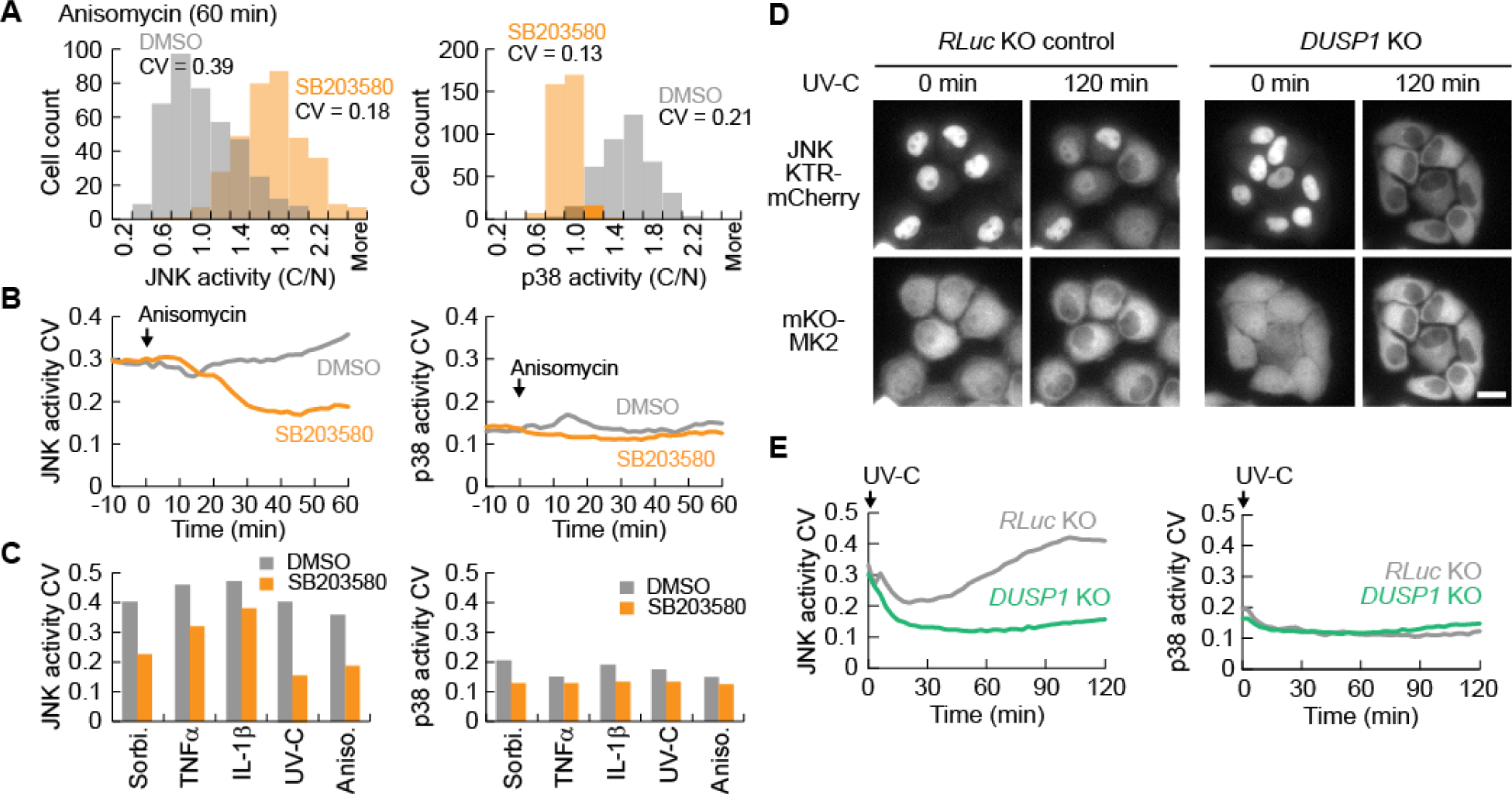
Cell-to-cell variability of stress and cytokine-induced JNK activities arises from cross-inhibition by p38. (**A**) Histograms of the JNK KTR-mCherry and mKO-MK2 C/N ratio distribution in 0.1% DMSO or 10 μM SB203580-pretreated cells at 60 min of 10 ng/ml anisomycin stimulation. The coefficient of variation (CV) is indicated. n ≥ 350 cells per condition from four independent experiments. (**B**) The time course of the CV upon 10 ng/ml anisomycin stimulation is shown for the JNK KTR-mCherry and mKO-MK2 C/N ratios in 0.1% DMSO or 10 μM SB203580-pretreated cells. n = 50 cells per condition from two independent experiments. (**C**) The CV of the JNK KTR-mCherry and mKO-MK2 C/N ratios at 30 min of 10 ng/ml TNFα, 60 min of 10 ng/ml IL-1β, 60 min of 10 ng/ml anisomycin, 60 min of 200 mM sorbitol, or 60 min of 100 J/m^2^ UV-C treatment are shown for 0.1% DMSO or 10 μM SB203580-pretreated clonal reporter cells. n = 50 cells per condition from two independent experiments. (**D**) HeLa *DUSP1* KO and *RLuc* KO control cells stably expressing NJP were stimulated with 100 J/m^2^ UV-C. Representative images of JNK KTR-mCherry and mKO-MK2 are shown. Scale bar: 20 μm. (**E**) The time course of the CV upon 100 J/m^2^ UV-C stimulation is shown for the JNK KTR-mCherry and mKO-MK2 C/N ratios in *DUSP1* KO and *RLuc* KO control cells. n = 50 cells per condition from two independent experiments.

### Correlation of JNK activity with fractional killing upon UV-C stress

Finally, we asked whether the cell-to-cell variability in JNK activity had an effect on cellular phenotypes. For this purpose, we studied UV-C-induced cell death, since it has been established that JNK activity is required for the induction of apoptosis in response to UV-C irradiation (*41, 42*). We exposed reporter cells to 100 J/m^2^ UV-C, and then tracked JNK and p38 activities and cellular outcomes, namely apoptosis or survival, in individual cells for 12 h (Fig. 4A). As has already been shown in Fig. 2, we observed an initial peak of JNK and p38 activities within 30 min, followed by downregulation, in both surviving and dying cells (Fig. 4A). Importantly, however, 2 h after UV-C irradiation JNK activity rose again and more prominently in the cells that were doomed to undergo apoptosis (Fig. 4, B and C). Meanwhile, almost all cells showed sustained p38 activation after the initial transient activation (Fig. 4, B and C).

**Fig. 4.**
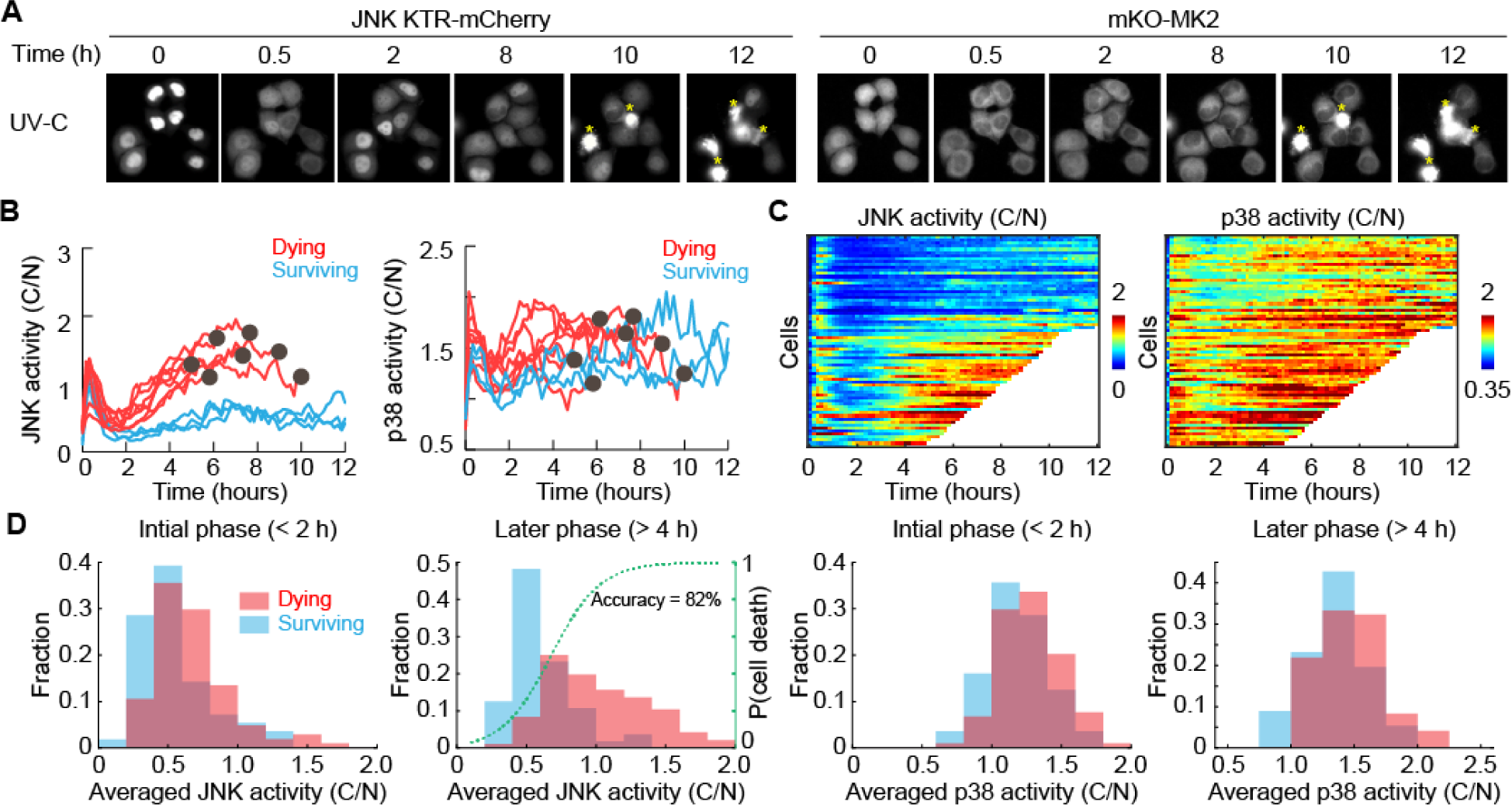
Cell-to-cell variation in JNK activity leads to fractional killing upon UV-C treatment. (**A**) Clonal HeLa cells stably expressing JNK KTR-mCherry, mKO-MK2, and NLS-iRFP-NLS were exposed to 100 J/m^2^ UV-C and imaged over time. Representative images are shown. Yellow asterisks indicate apoptotic cells. Scale bar: 20 μm. (**B**) C/N ratios of JNK KTR-mCherry and mKO-MK2 were quantified for dying (red lines) and surviving (blue lines) cells after 100 J/m^2^ UV-C stimulation. n = 10 representative cells. Black dots indicate apoptosis. (**C**) JNK KTR-mCherry and mKO-MK2 C/N ratios are displayed as heat maps for cells from (B). White traces indicate cells that underwent cell death. n = 70 cells from four independent experiments. (**D**) Histograms of average JNK and p38 activity in the initial phase (< 2 h) and later phase (> 4 h) of UV-C stimulation are shown in dying (red) and surviving (blue) cells. The green dashed line shows the estimated probability of cell death as a function of JNK activity, which is produced by the logistic regression.

To determine whether JNK and p38 activities were associated with cell death, we extracted the averaged JNK and p38 activities of single cells in the initial phase (< 2 h) and later phase (> 4 h), and compared these parameters between dying and surviving cells (Fig. 4D). We found that the mean JNK activities in the later phase were significantly associated with cell death, whereas the other three parameters were not significantly different between dying and surviving cells (Fig. 4D). We used a logistic regression to estimate the probability that a cell would die as a function of the mean JNK activity, resulting in a threshold of JNK activity (C/N) of 0.68 (Fig. 4D, green curve). Consistent with earlier reports (*36, 41, 42*), pretreatment with a JNK inhibitor significantly ameliorated UV-C-induced apoptosis (fig. S4A). Notably, the effect of adding a JNK inhibitor 2 h after UV-C irradiation was indistinguishable from the effect of pretreatment, indicating that the JNK activation in the later phase not only correlates with, but also promotes apoptosis (fig. S4B). SB203580 blocked UV-C-induced activation of p38 and resulted in sustained JNK activity (fig. S4, C and D), suggesting that p38-mediated repression of JNK activities caused the cell-to-cell-variability in JNK activities on longer time-scales as well. Although we hypothesized that SB203580 would increase or accelerate apoptosis following UV-C, the fraction of apoptotic cells was similar between SB203580- and DMSO-pretreated cells (fig. S4A). It is possible that p38 activation is also required for efficient induction of cell death; therefore we sought for a means of blocking the cross-inhibition of JNK without affecting p38 activity.

### Cell fate determination by variability of UV-C stress-induced DUSP induction

As we have already demonstrated (Fig. 2), DUSP1 is a major mediator of p38-mediated JNK cross-inhibition. Indeed, UV-C irradiation led to an increase in *DUSP1* mRNA levels in HeLa cells peaking at 2 h, which was completely blocked by SB203580 (fig. S4E). *DUSP1* KO cells exhibited enhanced and sustained JNK activities in all cells, while control cells exhibited heterogeneous responses (Fig. 5A). Increased JNK activities in *DUSP1* KO cells were accompanied by an increased rate of cell death (>90%) and a decreased period to cell death (Fig. 5, B and C). SB203580 treatment cancelled the effect of *DUSP1* deficiency in *DUSP1* KO cells (Fig. 5B), supporting the notion that *DUSP1* is the major p38-mediated suppressor of JNK-induced cell death. These data indicate that p38 plays roles in both pro-survival and pro-apoptotic pathways, and the former is mediated by cross-inhibition through p38-induced DUSP1 expression. Taken together, our data suggest that the fluctuation of p38-mediated DUSP1 induction efficiency leads to cell-to-cell heterogeneity in JNK activity and thereby in the cell fate in response to UV-C stress (Fig. 6).

**Fig. 5.**
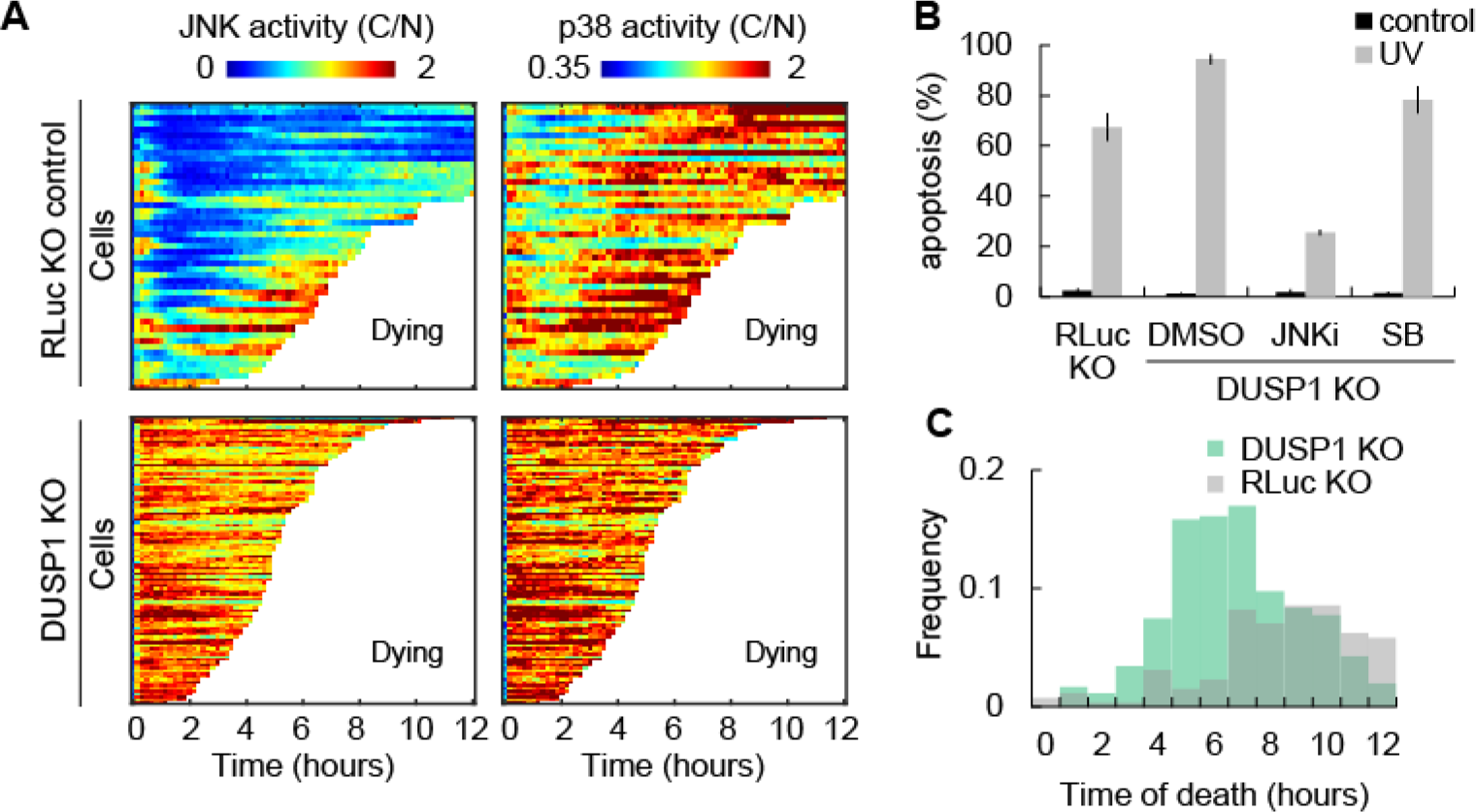
DUSP1 KO leads to loss of cell-to-cell heterogeneity of JNK activation and increase in cell death. (**A**) HeLa *DUSP1* KO and *RLuc* KO control cells stably expressing NJP were exposed to 100 J/m^2^ UV-C and imaged over time. JNK KTR-mCherry and mKO-MK2 C/N ratios are displayed as heat maps. n = 49 cells for *RLuc* KO and n = 100 cells for *DUSP1* KO. White traces indicate cells that underwent cell death. (**B**) Apoptosis of DMSO, 10 μM SB203580, or 10 μM JNK inhibitor VIII pretreated *DUSP1* KO cells and *RLuc* KO control cells 12 h after 100 J/m^2^ UV-C or mock treatment. Apoptosis was assayed by caspase 3 activation. Data are shown as the mean with SD. n = 3 independent experiments. (**C**) Histogram of the death time of *DUSP1* KO and *RLuc* KO HeLa cells upon 100 J/m^2^ UV-C treatment, determined by live-cell microscopy.

**Fig. 6.**
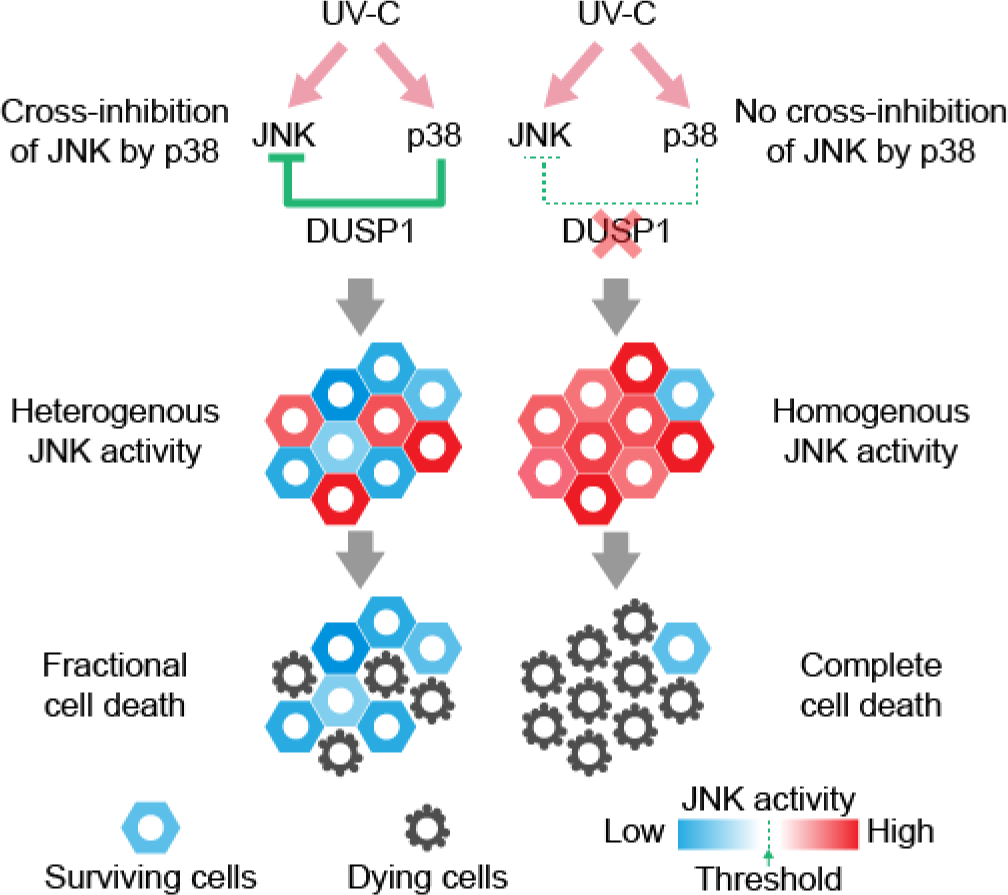
Cross-inhibition by p38 causes cell-to-cell variability of stress and cytokine-induced JNK activities. DUSP1-mediated cross-inhibition of JNK by p38 generates heterogeneity in JNK activity upon UV-C stress and leads to fractional killing (left). Depletion of DUSP1 reduces cell-to-cell variability of JNK activity, thereby inducing complete cell death (right).

## Discussion

In this study, we established a specific and sensitive imaging system for JNK and p38, which allowed us to quantitatively determine that p38 antagonizes JNK under diverse stress conditions by both post-translational and transcriptional mechanisms. Previous cell population-based studies have convincingly demonstrated that p38 negatively regulates JNK (*31–36*). While most of these studies identified single, predominant mechanisms that were responsible for this crosstalk in specific cell types and under specific conditions, our quantitative measurements enabled us to quantitate the contribution of post-translational and transcriptional cross-inhibition in each response to various inflammatory and stress stimuli (Fig. 2). Recently, Lauffenburger and his colleagues quantitatively demonstrated that p38 is a crucial factor for negative cross-talk to the JNK pathway under various conditions in synovial fibroblasts (*43*). They found that the compensatory effects of negative cross-talk evoked by inhibitors differ depending on the context of mitogenic or inflammatory stimulation. In our data, cross-inhibition by p38 generated cell-to-cell variability in JNK activity under all tested conditions, suggesting that this is a common mechanism underlying JNK and p38 signaling (Fig. 3C).

It has been well-established that JNK activity is required for UV-induced apoptosis by cell population-based studies (*41, 42*). We here found that UV-C stress induced heterogeneous JNK activation at the single cell level, and, strikingly, the cells with higher JNK activities underwent apoptosis, indicating the existence of a critical threshold of JNK activity in the cell-fate decision between apoptosis and survival (Fig. 4). Further, our data demonstrated that cross-inhibition by the p38-DUSP1 axis generates the cell-to-cell variability in JNK activity; thereby protecting the cells with low JNK activity from UV-C stress-induced apoptosis (Fig. 5 and 6). Consistently, Staples *et al.* demonstrated that UV-C induces *DUSP1* mRNA and protein in a p38-dependent manner and that DUSP1 protects cells from UV-C-induced apoptosis in MEFs by inactivating JNK (*36*). One possible mechanism underlying the cell-to-cell variability of JNK activity is likely to be the cell-to-cell variability of DUSP1 induction. For instance, intrinsic noise of gene expression results in an increase in cellular heterogeneity that is known as non-genetic heterogeneity, which eventually leads to resistance or persistence to anticancer drugs or antibiotics at the population level (*12, 13*). According to this mechanism, it is plausible that the heterogeneity of JNK activity generated by cross-inhibition from p38 confers resistance to cells against stress stimuli. On the other hand, it is largely unclear how heterogeneity in JNK activity is generated through post-translational cross-inhibition by p38, and is connected to physiological outcomes. Future studies will be needed to unveil the degree to which the cross-inhibition via p38 contributes to the cellular heterogeneity of JNK activity, and the mechanism underlying this effect.

As to the post-translational cross-inhibition mechanism, Cheung *et al.* proposed that p38 mediates negative feedback control of TAK1 and downstream kinases in response to pro-inflammatory cytokines and osmotic stress by p38-mediated phosphorylation of TAB1 on Ser423 and Thr431 (*31*). Our results strongly support this model, because the rescue experiment with the non-phosphorylatable TAB1 mutant further increased JNK activity upon stimulation with TNFα and IL-1β (fig. S2G). Interestingly, TAB1 phosphorylation did not affect p38 activity (fig. S2G). One possible explanation for this finding is that TAB1 is essential for TAK1 activity towards JNK, but not p38 in HeLa cells. This is supported by the observation that TNFα-induced JNK activity was fully impaired in *TAB1* and *TAK1* KO cells, whereas TNFα-induced p38 activity was strongly decreased in *TAK1* KO, but not changed in *TAB1* KO cells. Another possible mechanism is that a different p38-specific MAPKKK may compensate for the p38 activation. In fact, it has been reported that thousand-and-one amino acid kinase 2 (TAO2), a MAPKKK that preferentially activates the p38 pathway, is activated by sorbitol-induced osmotic stress (*44*). Meanwhile, there is no doubt that DUSP1 was involved in the transcriptional cross-inhibition of JNK, since it is known that p38 can induce *DUSP1* gene expression in response to pro-inflammatory cytokines and various stresses (fig. S2C and S4E) (*33, 36, 40, 45*). However, as with the case of cross-inhibition by post-translational modification, it is unclear why DUSP1, a known phosphatase for JNK and p38, would inactivate only a small part of p38 in our system, when cells were exposed to anisomycin and UV-C (Fig. 2E and fig. S2E). This observation is likely due to the substrate preference of DUSP1 for JNK over p38 (*46*). In line with our data, a previous study demonstrated that UV-C-stimulated JNK phosphorylation is quickly downregulated in MEFs within 90 min, concomitant with DUSP1 induction, while p38 phosphorylation decreases more slowly, with some level remaining up to 6 h (*36*).

Our imaging system is based on KTRs, which enabled simultaneous visualization of ERK, JNK, and p38 activities in single cells (*22*). Here, we presented a new live-cell imaging reporter for p38 activity based on the naturally shuttling p38 substrate MK2. Exogenous expression of MK2 did not show an apparent effect on the phenotype of HeLa cells during their maintenance; however, it remains to be determined whether the overexpression of MK2 perturbs the signaling and has an effect on the phenotype of cells of interest. Nonetheless, our highly specific and sensitive multiplexed imaging system will be useful to quantitatively evaluate the interplay of p38 and JNK, revealing the heterogeneity of their interaction among single cells, and ultimately the role of their activity dynamics in phenotypic outcomes in response to stress and inflammatory signals.

## Materials and Methods

### Plasmids

Plasmids were constructed by standard molecular biology methods. The NLS of SV40 large T antigen (PKKKRKV) was added to the N- and C-termini of iRFP by PCR and the construct was cloned into the pT2Apuro vector (*47*), a Tol2 transposon vector with IRES-*pac* (puromycin resistance gene), generating pT2Apuro-NLS-iRFP-NLS. The pT2A vector was a kind gift from Dr. K. Kawakami (National Institute for Genetics, Japan). JNK KTR was amplified by PCR from pLentiPGK Puro DEST JNKKTRClover (*22*), which was a kind gift from Dr. M. Covert (Addgene plasmid #59151), and cloned into the eukaryotic expression vector pCAGGS (*48*) or pPBbsr, a piggyBac transposon vector with IRES-*bsr* (blasticidin S resistance gene) (*49*), along with mCherry, resulting in pCAGGS-JNK KTR-mCherry and pPBbsr-JNK KTR-mCherry. The pPB backbone was a kind gift from Dr. A. Bradley (Wellcome Trust Sanger Institute, UK). p38 KTR (*22*) was synthesized by PCR from annealed sense- and antisense oligonucleotides and cloned into a pPBbsr-mEGFP vector to generate pPBbsr-p38 KTR-mEGFP. For pCAGGS- mKO-MK2, pPBbsr-mKO-MK2, and pPBbsr-mEGFP-MK2, the cDNA of human *MAPKAPK2* was amplified by PCR from a HeLa cDNA library and inserted into the pCAGGS or pPBbsr backbone together with mEGFP or mKO. To construct a polycistronic vector encoding NLS-iRFP-NLS, JNK KTR-mCherry, and mKO-MK2, sequences for the self-cleaving P2A peptide (GSGATNFSLLKQAGDVEENPGP) were PCR-cloned between the reporters, and the polycistronic cassette was inserted into the pPBbsr backbone to generate pNJP. The cDNA of human TAB1 was obtained from HeLa cDNA by PCR, and the S423A and T431A point mutations were introduced into TAB1 by two-step overlap-extension PCR. The cDNAs of TAB1-wt and SATA were subcloned into pCSIIneo vector, a lentivirus vector with IRES-*neo* (neomycin resistance gene), to generate pCSIIneo-TAB1-wt and pCSIIneo-TAB1-SATA. The lentiviral CSII-EF backbone vector was a kind gift from Dr. H. Miyoshi (RIKEN, Japan). lentiCRISPR v2 was a kind gift from Dr. F. Zhang (Addgene plasmid #52961) (*50*).

### Reagents

Anisomycin and actinomycin D were purchased from Sigma Aldrich, JNK inhibitor VIII from Calbiochem, SB203580 from Selleck Chemicals, and VX-745 from AdipoGen. Sorbitol was from Wako. Recombinant human TNFα and recombinant human IL-1β were acquired from R&D Systems. Blasticidin S, puromycin, and G418 were purchased from InvivoGen. The primary antibodies against p38 (#9212), phospho-p38 (Thr180/Tyr182) (#9216), JNK (#9252), phospho-JNK (Thr183/Tyr185) (#9355), JNK1 (#3708), JNK2 (#9258), TAK1 (#5206) and TAB1 (#3226) were purchased from Cell Signaling Technology. Anti-α-tubulin (CP06) was obtained from Calbiochem. Anti-mCherry (ab167453) and anti-monomeric Kusabira Orange 2 (PM051M) antibodies were obtained from Abcam and MBL, respectively. Secondary antibodies IRDye 680LT goat anti-rabbit IgG and IRDye 800CW donkey anti-mouse IgG were purchased from LI-COR Biosciences. Irradiation with a defined dose of 100 J/m^2^ UV-C light (254 nm) was performed using a CL-100 Ultraviolet Crosslinker (UVP), while most of the medium was removed. In a four-compartment 35 mm glass-bottom dish, the cells were irradiated, then covered with 100 μl imaging medium per well to keep them from drying out.

### Cell culture and cell line generation

HeLa cells were purchased from the Human Science Research Resources Bank (Japan) and were maintained in Dulbecco’s Modified Eagle’s Medium (DMEM) high glucose (Wako; Nacalai) supplemented with 10% fetal bovine serum (Sigma Aldrich). Antibiotics were added, when applicable. For transient expression, pCAGGS vectors were transfected using 293fectin (Invitrogen) according to the manufacturer’s instructions. To establish stable cell lines expressing the fluorescent reporters, PiggyBac- and Tol2 transposon-based systems were applied. The reporters encoding pPBbsr or pT2Apuro vectors were co-transfected with the transposase encoding pCMV-mPBase (neo-) vector (a kind gift of Dr. A. Bradley, Wellcome Trust Sanger Institute, UK) and pCAGGS-T2TP vector (a kind gift of Dr. K. Kawakami, National Institute for Genetics, Japan), respectively, using 293fectin. One day after transfection, cells were selected with 20 μg/ml blasticidin S or 1 μg/ml puromycin for at least one week. To obtain the triple reporter cell line expressing NLS-iRFP-NLS, JNK KTR-mCherry, and mKO-MK2 simultaneously, the reporters were introduced based on the PiggyBac and Tol2 transposon system, and single cells were cloned and screened for iRFP, mCherry, and mKO expression. For easier generation of KO cell lines expressing the three reporters, the polycistronic vector pNJP was stably transfected using the PiggyBac transposon system. However, the expression levels of the polycistronic construct were reduced compared to the individually expressed reporters (Fig. S1I). To complement *TAB1* KO cells with the wild-type or mutant TAB1, lentivirus-mediated gene transfer was used. In brief, the lentiviral pCSIIneo vector was transfected into Lenti-X 293T cells (Clontech) together with the packaging plasmid psPAX2, which was a gift from Dr. D. Trono (Addgene plasmid #12260), and pCMV-VSV-G-RSV-Rev (a kind gift of Dr. Miyoshi, RIKEN, Japan) by using the linear polyethyleneimine “Max” MW 40,000 (Polyscience). After two days, target cells were infected with the virus-containing media. Beginning at two days after infection, the cells were selected by at least one week of treatment with 1 mg/ml G418. All cells were maintained in a humidified atmosphere of 5% CO_2_ at 37 °C.

### CRISPR/Cas9-mediated KO

For CRISPR/Cas9-mediated KO of human *DUSP1*, *MAPK8*, *MAPK9*, *MAPK14*, *MAP3K7*, and *TAB1* genes, single guide RNAs (sgRNA) targeting the start codon or first exon were designed using the CRISPR Design online tool (http://crispr.mit.edu, Zhang Lab, MIT). The *TAB1* sgRNA targeted an upstream sequence of the start codon, allowing overexpression of TAB1 in *TAB1* KO cells which stably express the sgRNA and Cas9. The following targeting sequences were used: *DUSP1*, CTACTAACCTGATCGTAGAG; *MAPK8*, ACGCTTGCTTCTGCTCATGA; *MAPK9*, TCAGTTTTATAGTGTGCAAG; *MAPK14*, AGCTCCTGCCGGTAGAACGT; *MAP3K7*, CATCTCACCGGCCGAAGACG; *RLuc8*, AGGTGTACGACCCCGAGCAG; *TAB1*,CCTCCTCTGCGCCGCCATCT. Annealed Oligo DNAs for the sgRNAs were cloned into the lentiCRISPR v2 vector (Addgene plasmid #52961) (*50*) and the sgRNA/Cas9 cassettes were introduced into HeLa cells by lentiviral gene transfer. Infected cells were selected using 2 μg/ml puromycin and single cells were cloned. KO was confirmed by western blotting or the TIDE webtool (https://tide-calculator.nki.nl/) (*51*).

### Western blotting

HeLa cells were starved in DMEM medium for at least 3 h, unless otherwise specified, treated with inhibitors and stimulants, when indicated, and lysed in 1x SDS sample buffer (62.5 mM Tris-HCl pH 6.8, 12% glycerol, 2% SDS, 0.004% Bromo Phenol Blue, and 5% 2-mercaptoethanol). After sonication, the samples were separated by SDS-polyacrylamide gel electrophoresis and transferred to polyvinylidene membranes (Millipore). After blocking with Odyssey blocking buffer (LI-COR) or skim milk for 1 h, the membranes were incubated with primary antibodies diluted in Odyssey blocking buffer or TBS Tween20 overnight, followed by secondary antibodies diluted in Odyssey blocking buffer. Fluorescent signals were detected by an Odyssey Infrared scanner (LI-COR) and analyzed by the Odyssey imaging software.

### RNA preparation and quantitative PCR analysis

RNA was isolated with a QIAshredder Kit (QIAGEN) and RNeasy Plus Mini Kit (QIAGEN), according to the manufacturer’s instructions. The extracted total RNA was reverse-transcribed using the High-Capacity cDNA Reverse Transcription Kit (Thermo Fisher Scientific).

Quantitative PCR analyses were performed with the Power SYBR Green PCR Master Mix (Thermo Fisher Scientific) or THUNDERBIRD SYBR qPCR Mix (TOYOBO) on an Applied Biosystems StepOne Real-Time PCR System (Thermo Fisher Scientific). The ΔΔCt method was used to normalize data to the housekeeping gene *GAPDH* and to present data as fold changes compared to control samples (*52*). Primer sequences were as follows: *DUSP1* Fw: ACCACCACCGTGTTCAACTTC; *DUSP1* Rv: TGGGAGAGGTCGTAATGGGG; *GAPDH* Fw: GAGTCCACTGGCGTCTTCAC; *GAPDH* Rv: GTTCACACCCATGACGAACA.

### Live cell imaging

HeLa cells stably expressing the fluorescent reporters were grown on 35 mm glass-bottom dishes (Asahi Techno Glass) or four-compartment 35 mm glass-bottom dishes (Greiner Bio-One) for at least 24 h and starved in FluoroBrite DMEM (Life Technologies) supplemented with 1x GlutaMax (Life Technologies) and 0.2% fetal bovine serum for about 3 h. When applicable, cells were pretreated with inhibitors for about 15 min before imaging and then treated with stimulants. For short-term imaging experiments, images were acquired on an IX81 inverted microscope (Olympus), which was equipped with a Retiga 4000R cooled Mono CCD camera (QImaging), a Spectra-X light engine illumination system (Lumencor), an IX2-ZDC laser-based autofocusing system (Olympus), a UPlanSApo 60x/1.35 oil objective lens (Olympus), a MAC5000 controller for filter wheels and XY stage (Ludl Electronic Products), an incubation chamber (Tokai Hit), and a GM-4000 CO_2_ supplier (Tokai Hit). The following filters and dichroic mirrors were used: for mKO, an FF01-543/3 excitation filter (Semrock), a 20/80 beamsplitter dichroic mirror (Chroma), and an FF01-563/9 emission filter (Semrock); for mCherry, an FF01-580/20 excitation filter (Semrock), a 20/80 beamsplitter dichroic mirror (Chroma), and an FF01-641/75 emission filter (Semrock); for iRFP, an FF02-632/22 excitation filter (Semrock), an FF408/504/581/667/762-Di01 dichroic mirror (Semrock), and an FF01-692/LP emission filter (Semrock). For long-term imaging, an IX83 inverted microscope (Olympus) was used, which was equipped with a Prime sCMOS camera (Photometrix), a Spectra-X light engine illumination system (Lumencor), an IX3-ZDC2 laser-based autofocusing system (Olympus), UPlanSApo 20x/0.75 and UPlanSApo 40x2/0.75 dry objective lenses (Olympus), a U-CBF control box for filter wheels and IX3-SSU ultrasonic stage (Olympus), an IX3WX incubation chamber (Tokai Hit), and an STX Stage Top Incubator (Tokai Hit). A BK7 glass dichroic mirror (EKSMA Optics) and the following filters were used: for iRFP, a BLP01-664R emission filter (Semrock); for mCherry, an FF01-575/15 excitation filter (Semrock) and an FF02-641/75 emission filter (Semrock); for mKO, an FF01-543/3 excitation filter (Semrock) and an FF01-563/9 emission filter (Semrock). For FRET imaging on the IX81 inverted microscope, an XF2034 455DRLP dichroic mirror (Omega Optical) and the following filters were used: for FRET, an FF01-542/27 emission filter (Semrock); for CFP, an FF01-483/32 emission filter (Semrock). The microscopes were controlled by MetaMorph software (Molecular Devices).

### Image analysis

Images were processed using the MetaMorph software as previously described (*53*). For multiplexed imaging of translocation reporters, the background was corrected by subtraction of blank positions (medium only) or minimum planes to flat-field the images. Regions of interest were set in the nucleus and cytosol of single cells and the average value of their fluorescence intensities was measured. The ratio of the cytosolic and nuclear intensity (C/N ratio) was calculated using Excel software (Microsoft Corporation).

### Quantification of signaling activities

The strength of signaling activities was quantified in Fig. 2D as follows: First, C/N ratios of JNK-KTR at the indicated time point for each stimulation were picked up in the presence of SB203580, and these values were assigned as the stimulation-to-JNK edge activities. In a similar way, each stimulation-to-p38 edge activity was retrieved from the C/N ratio of mKO-MK2 under a DMSO-treated condition. Next, the post-translational cross-inhibition activity by p38 for each stimulation was calculated by subtracting the C/N ratio of JNK-KTR under an actinomycin D-treated condition from that under an SB203580-treated condition at the indicated time point.

Finally, the transcriptional cross-inhibition activity by p38 was calculated by subtracting the C/N ratio of JNK-KTR under a DMSO-treated condition from that under an actinomycin D-treated condition.

### Apoptosis assay

HeLa cells stably expressing the caspase 3 FRET biosensor ECRP (*54*) were seeded to four-compartment 35 mm glass-bottom dishes (Greiner Bio-One) at a density of 3 x 10^4^ cells/well. One day after seeding, cells were starved in FluoroBrite DMEM (Life Technologies) supplemented with 1x GlutaMax (Life Technologies) and 0.2% fetal bovine serum for 3 h, followed by inhibitor pretreatment for about 15 min and then UV-C stimulation, when indicated. At 12 h after UV-C, the cells were imaged by an IX81 inverted microscope with a 20x objective lens, as described earlier. FRET and CFP images were obtained in 4 to 5 positions per well and at least 3 independent experiments were performed. After subtraction of a minimum projection image to flat-field the images, the FRET and CFP intensities were measured for single cells and the FRET/CFP ratio was calculated in Excel software. A minimum of 2000 cells were analyzed per condition. Cells showed a bimodal FRET/CFP ratio distribution and cells with a low FRET/CFP ratio (≤ 2), corresponding to high caspase 3 activity, were counted as apoptotic. Time-lapse imaging confirmed that within 12 h post UV-C, cells did not detach from the plate.

### Regression analysis

For non-linear regression of dose responses, we utilized solver functions in Microsoft Excel and fitted the experimental data with the Hill function to obtain EC50 and nH values. Logistic regression analysis was performed using Python.

### Statistical analysis

Statistical analysis was conducted using Microsoft Excel software. For comparison of two datasets, a two tailed, unpaired Student’s *t*-test was performed according to the result of the F-test. *p-*values < 0.05 were considered significant at the following levels: \**p* < 0.05, \*\**p* < 0.01, \*\*\**p* < 0.001.

## Supplementary Materials

Fig. S1. Characterization of a multiplexed imaging system for JNK and p38 kinase activities.

Fig. S2. Cross-inhibition of JNK by p38.

Fig. S3. Cell-to-cell variability of JNK and p38 activity upon inflammatory and stress stimuli. Fig. S4. Cell-to-cell variation in JNK activity leads to fractional killing upon UV-C treatment. Movie S1. Multiplexed imaging of JNK and p38 activities upon anisomycin treatment.

Movie S2. Multiplexed imaging of JNK and p38 activities upon UV-C stress.

## Acknowledgments

We thank Dr. A. Bradley (pPB, pCMV-mPBase), Dr. M. Covert (pLentiPGK Puro DEST JNKKTRClover), Dr. K. Kawakami (pT2A, pCAGGS-T2TP), Dr. H. Miyoshi (CSII-EF, pCMV-VSV-G-RSV-Rev), Dr. D. Trono (psPAX2), and Dr. F. Zhang (lentiCRISPR v2) for providing plasmids. We are grateful to Dr. T. Tomida for providing a plasmid (perky p38 NES) and for helpful discussions. We also thank the members of the Matsuda Laboratory and Aoki Laboratory for their productive discussions. A. Kawagishi, N. Sakikawa, K. Hirano, N. Nishimoto, and E. Ebine are to be thanked for their technical assistance.

## Author contributions

H.M. and K.A. designed the research; H.M. performed the experiments. H.M. and Y.K. analyzed the data. H.M., K.A. and M.M. wrote the paper.

## References and Notes

1. B. G. Childs, D. J. Baker, J. L. Kirkland, J. Campisi, J. M. van Deursen, Senescence and apoptosis: dueling or complementary cell fates? EMBO Rep. 15, 1139–53 (2014).

2. K. H. Vousden, D. P. Lane, p53 in health and disease. Nat. Rev. Mol. Cell Biol. 8, 275 (2007).

3. A. J. Levine, p53, the cellular gatekeeper for growth and division. Cell. 88, 323–31 (1997).

4. X. Chen, L. J. Ko, L. Jayaraman, C. Prives, p53 levels, functional domains, and DNA damage determine the extent of the apoptotic response of tumor cells. Genes Dev. 10, 2438–51 (1996).

5. M. Kracikova, G. Akiri, A. George, R. Sachidanandam, S. A. Aaronson, A threshold mechanism mediates p53 cell fate decision between growth arrest and apoptosis. Cell Death Differ. 20, 576–588 (2013).

6. C. J. Marshall, Specificity of receptor tyrosine kinase signaling: transient versus sustained extracellular signal-regulated kinase activation. Cell. 80, 179–85 (1995).

7. S. L. Werner, D. Barken, A. Hoffmann, Stimulus specificity of gene expression programs determined by temporal control of IKK activity. Science. 309, 1857–61 (2005).

8. J. E. Purvis et al., P53 Dynamics Control Cell Fate. Science. 336, 1440–4 (2012).

9. J. E. Purvis, G. Lahav, Encoding and decoding cellular information through signaling dynamics. Cell. 152, 945–956 (2013).

10. K. Takemoto, T. Nagai, A. Miyawaki, M. Miura, Spatio-temporal activation of caspase revealed by indicator that is insensitive to environmental effects. J. Cell Biol. 160, 235–43 (2003).

11. C. Holohan, S. Van Schaeybroeck, D. B. Longley, P. G. Johnston, Cancer drug resistance: an evolving paradigm. Nat. Rev. Cancer. 13, 714–26 (2013).

12. A. Brock, H. Chang, S. Huang, Non-genetic heterogeneity--a mutation-independent driving force for the somatic evolution of tumours. Nat. Rev. Genet. 10, 336–42 (2009).

13. Y. Wakamoto et al., Dynamic Persistence of Antibiotic-Stressed Mycobacteria. Science (80-.). 339, 91–95 (2013).

14. S. L. Spencer, S. Gaudet, J. G. Albeck, J. M. Burke, P. K. Sorger, Non-genetic origins of cell-to-cell variability in TRAIL-induced apoptosis. Nature. 459, 428–432 (2009).

15. J. Roux et al., Fractional killing arises from cell-to-cell variability in overcoming a caspase activity threshold. Mol. Syst. Biol. 11, 1–18 (2015).

16. A. L. Paek et al., Cell-to-Cell Variation in p53 Dynamics Leads to Fractional Killing. Cell. 165, 1–12 (2016).

17. G. L. Johnson, R. Lapadat, Mitogen-Activated Protein Kinase Pathways Mediated by ERK, JNK, and p38 Protein Kinases The Protein Kinase Complement of the Human Genome. Science. 298, 1911–1912 (2002).

18. A. Cuadrado, A. R. Nebreda, Mechanisms and functions of p38 MAPK signalling. Biochem. J. 429, 403–417 (2010).

19. C. R. Weston, R. J. Davis, The JNK signal transduction pathway. Curr. Opin. Cell Biol. 19, 142–149 (2007).

20. E. F. Wagner, Á. R. Nebreda, Signal integration by JNK and p38 MAPK pathways in cancer development. Nat. Rev. Cancer. 9, 537–549 (2009).

21. M. Raman, W. Chen, M. H. Cobb, Differential regulation and properties of MAPKs. Oncogene. 26, 3100–3112 (2007).

22. S. Regot, J. J. Hughey, B. T. Bajar, S. Carrasco, M. W. Covert, High-sensitivity measurements of multiple kinase activities in live single cells. Cell. 157, 1724–34 (2014).

23. N. Hao, B. A. Budnik, J. Gunawardena, E. K. O’Shea, Tunable signal processing through modular control of transcription factor translocation. Science. 339, 460–4 (2013).

24. A. Komeili, E. K. O’Shea, Roles of phosphorylation sites in regulating activity of the transcription factor Pho4. Science (80-.). 284, 977–980 (1999).

25. J. D. Nardozzi, K. Lott, G. Cingolani, Phosphorylation meets nuclear import: a review. Cell Commun. Signal. 8, 32 (2010).

26. T. Obsil, V. Obsilova, Structure/function relationships underlying regulation of FOXO transcription factors. Oncogene. 27, 2263–2275 (2008).

27. K. Engel, A. Kotlyarov, M. Gaestel, Leptomycin B-sensitive nuclear export of MAPKAP kinase 2 is regulated by phosphorylation. EMBO J. 17, 3363–71 (1998).

28. W. Meng et al., Structure of mitogen-activated protein kinase-activated protein (MAPKAP) kinase 2 suggests a bifunctional switch that couples kinase activation with nuclear export. J. Biol. Chem. 277, 37401–37405 (2002).

29. O. J. Trask et al., Assay Development and Case History of a 32K-Biased Library High-Content MK2-EGFP Translocation Screen to Identify p38 Mitogen-Activated Protein Kinase Inhibitors on the ArrayScan 3.1 Imaging Platform. Methods Enzymol. 414, 419– 439 (2006).

30. J. H. Kim et al., High cleavage efficiency of a 2A peptide derived from porcine teschovirus-1 in human cell lines, zebrafish and mice. PLoS One. 6, 1–8 (2011).

31. P. C. F. Cheung, D. G. Campbell, A. R. Nebreda, P. Cohen, Feedback control of the protein kinase TAK1 by SAPK2a/p38alpha. EMBO J. 22, 5793–805 (2003).

32. L. Hui et al., p38alpha suppresses normal and cancer cell proliferation by antagonizing the JNK-c-Jun pathway. Nat. Genet. 39, 741–9 (2007).

33. C. Kim et al., The kinase p38 alpha serves cell type-specific inflammatory functions in skin injury and coordinates pro- and anti-inflammatory gene expression. Nat. Immunol. 9, 1019–27 (2008).

34. J. Heinrichsdorff, T. Luedde, E. Perdiguero, A. R. Nebreda, M. Pasparakis, p38 alpha MAPK inhibits JNK activation and collaborates with IkappaB kinase 2 to prevent endotoxin-induced liver failure. EMBO Rep. 9, 1048–54 (2008).

35. C. Caballero-Franco et al., Tuning of protein kinase circuitry by p38α is vital for epithelial tissue homeostasis. J. Biol. Chem. 288, 23788–97 (2013).

36. C. J. Staples, D. M. Owens, J. V. Maier, A. C. B. Cato, S. M. Keyse, Cross-talk between the p38α and JNK MAPK pathways mediated by MAP kinase phosphatase-1 determines cellular sensitivity to UV radiation. J. Biol. Chem. 285, 25928–25940 (2010).

37. L. Pereira, A. Igea, B. Canovas, I. Dolado, A. R. Nebreda, Inhibition of p38 MAPK sensitizes tumour cells to cisplatin-induced apoptosis mediated by reactive oxygen species and JNK. EMBO Mol. Med. 5, 1759–1774 (2013).

38. E. Perdiguero et al., Genetic analysis of p38 MAP kinases in myogenesis: fundamental role of p38alpha in abrogating myoblast proliferation. Embo J. 26, 1245–1256 (2007).

39. J. S. C. Arthur, S. C. Ley, Mitogen-activated protein kinases in innate immunity. Nat. Rev. Immunol. 13, 679–692 (2013).

40. I. Ferreiro et al., Whole genome analysis of p38 SAPK-mediated gene expression upon stress. BMC Genomics. 11, 144 (2010).

41. Y. Chen, X. Wang, D. Templeton, R. J. Davis, T. Tan, The Role of c-Jun N-terminal Kinase (JNK) in Apoptosis Induced by Ultraviolet C and Y Radiation. J. Biol. Chem. 271, 31929–31936 (1996).

42. C. Tournier et al., Requirement of JNK for stress-induced activation of the cytochrome c-mediated death pathway. Science. 288, 870–4 (2000).

43. D. S. Jones, A. P. Jenney, B. A. Joughin, P. K. Sorger, D. A. Lauffenburger, Sci. Signal., in press, doi:10.1126/scisignal.aal1601.

44. Z. Chen, M. H. Cobb, Regulation of Stress-responsive Mitogen-activated Protein (MAP) Kinase Pathways by TAO2. J. Biol. Chem. 276, 16070–16075 (2001).

45. T. Tomida, M. Takekawa, H. Saito, Oscillation of p38 activity controls efficient pro-inflammatory gene expression. Nat. Commun. 6, 8350 (2015).

46. D. N. Slack, O. M. Seternes, M. Gabrielsen, S. M. Keyse, Distinct binding determinants for ERK2/p38alpha and JNK map kinases mediate catalytic activation and substrate selectivity of map kinase phosphatase-1. J. Biol. Chem. 276, 16491–500 (2001).

47. K. Kawakami, T. Noda, Transposition of the Tol2 Element, an Ac-Like Element from the Japanese Medaka Fish Oryzias latipes, in Mouse Embryonic Stem Cells. Genetics. 166, 895–899 (2004).

48. H. Niwa, K. Yamamura, J. Miyazaki, Efficient selection for high-expression transfectants with a novel eukaryotic vector. Gene. 108, 193–9 (1991).

49. K. Yusa, R. Rad, J. Takeda, A. Bradley, Generation of transgene-free induced pluripotent mouse stem cells by the piggyBac transposon. Nat. Methods. 6, 363–369 (2009).

50. N. E. Sanjana, O. Shalem, F. Zhang, Improved vectors and genome-wide libraries for CRISPR screening. Nat. Methods. 11, 783–784 (2014).

51. E. K. Brinkman, T. Chen, M. Amendola, B. Van Steensel, Easy quantitative assessment of genome editing by sequence trace decomposition. Nucleic Acids Res. 42, 1–8 (2014).

52. T. D. Schmittgen, K. J. Livak, Analyzing real-time PCR data by the comparative CT method. Nat. Protoc. 3, 1101–1108 (2008).

53. G. Maryu, M. Matsuda, K. Aoki, Multiplexed fluorescence imaging of ERK and Akt activities and cell-cycle progression. Cell Struct. Funct. 92, 81–92 (2016).

54. J. G. Albeck et al., Quantitative Analysis of Pathways Controlling Extrinsic Apoptosis in Single Cells. Mol. Cell. 30, 11–25 (2008).

