## Supplementary Materials for "Cell-to-cell heterogeneity in p38-mediated cross-inhibition of JNK causes stochastic cell death"

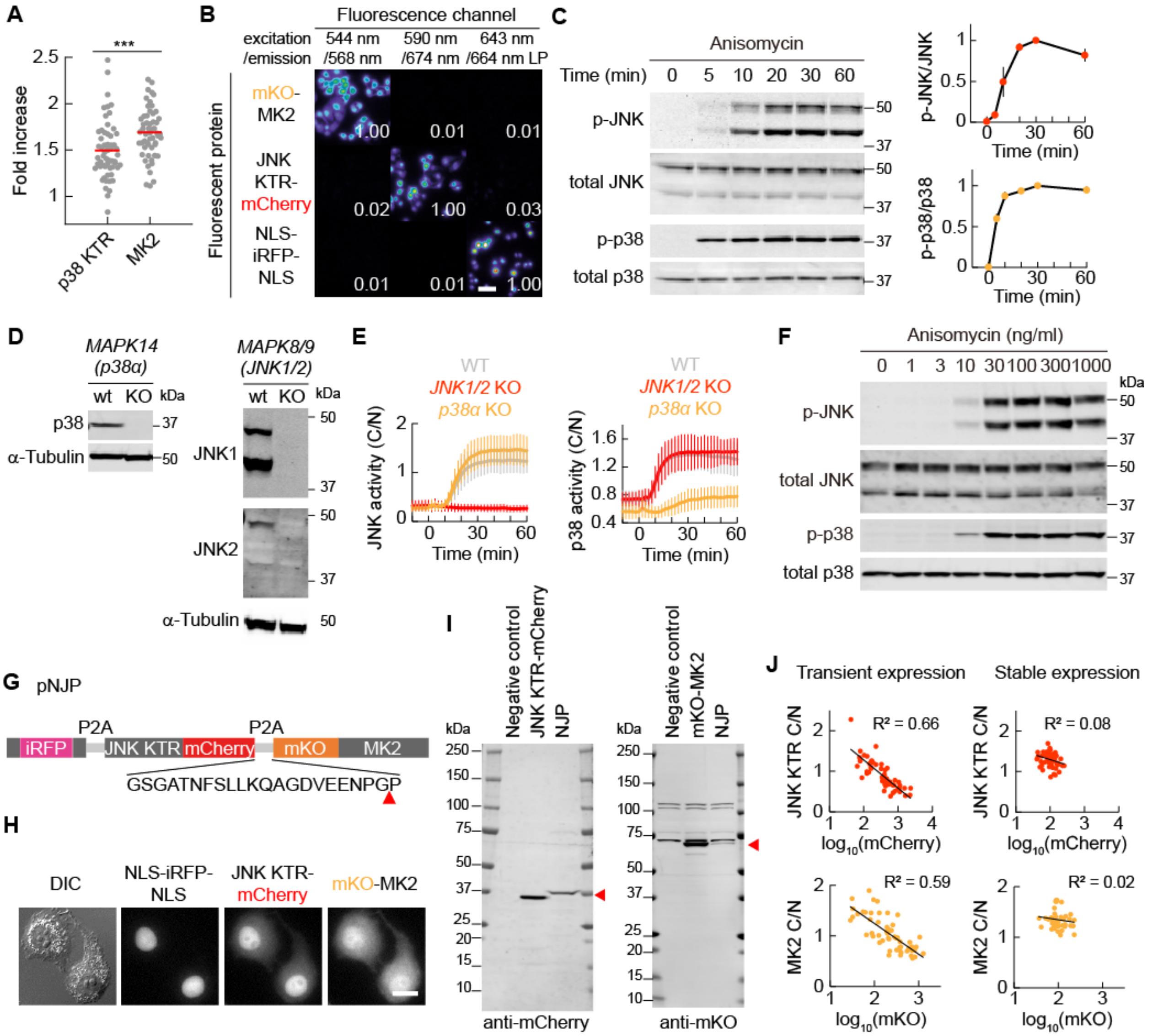


**Fig. S1. Characterization of a multiplexed imaging system for JNK and p38 kinase activities.** (**A**) The fold increases in the C/N ratios of p38 KTR-mEGFP and mEGFP-MK2 after 30 min of 1 μg/ml anisomycin stimulation are shown for single cells and as an average (red line). n ≥ 50 cells for each condition from two independent experiments. Significance was tested by a Student's *t*-test. ***p<0.001. (**B**) Bleed-through analysis of JNK KTR-mCherry, mKO-MK2, and NLS-iRFP-NLS. HeLa cells stably expressing single reporters were imaged in each fluorescent channel and images are represented in a pseudo-color scale. The values shown represent the fraction of fluorescence leakage. Scale bar: 60 μm. (**C**) HeLa cells were treated with 1 μg/ml anisomycin for the indicated times. Cell lysates were analyzed by western blotting against p-p38, total p38, p-JNK, and total JNK. Representative blots out of three independent experiments are shown (left). The time courses of the p-JNK/JNK and p-p38/p38 ratios upon 1 μg/ml anisomycin stimulation are plotted (right). All time series data were normalized between 0 and 1 using minimum and maximum values for comparison. Data are presented as the mean with SD. n = 3 independent experiments. (**D**) KO of *MAPK14* (*p38α*) and double KO of *MAPK8* (*JNK1*) and *MAPK9* (*JNK2*) were confirmed by western blotting against total p38, JNK1, JNK2, and α-tubulin as a control. (**E**) HeLa wildtype, *MAPK8/9* (*JNK1/2*) double KO, and *MAPK14* (*p38α*) KO cells stably expressing NJP were stimulated with 1 μg/ml anisomycin. Averaged JNK KTR-mCherry and mKO-MK2 C/N ratios with SD are shown. n = 20 cells for each condition. (**F**) HeLa cells were treated with the indicated doses of anisomycin for 1 h. Cell lysates were analyzed by western blotting against p-p38, total p38, p-JNK, and total JNK. Representative blots out of three independent experiments are shown. (**G**) Plasmid structure of the polycistronic pNJP reporter construct, consisting of NLS-iRFP-NLS, JNK KTR-mCherry, and mKO-MK2, connected by self-cleaving P2A peptide sequences, so that the reporters are expressed separately. The P2A peptide sequence and point of cleavage (red arrowhead) are indicated. (**H**) Representative images of NLS-iRFP-NLS, JNK KTR-mCherry, and mKO-MK2 stably expressed from pNJP in HeLa cells are shown. Scale bars: 20 μm. (**I**) HeLa control cells and cells stably expressing JNK KTR-mCherrry, mKO-MK2 or the polycistronic construct NJP were lysed and analyzed by western blotting against mCherry and mKO. JNK KTR-mCherry (33.6 kDa) and mKO-MK2 (70.2 kDa) samples served as a positive control (red arrowhead). (**J**) JNK KTR-mCherry and mKO-MK2 were transiently overexpressed (left) or stably expressed by transposon-mediated gene-transfer, i.e., the PiggyBac transposon system (right) in HeLa cells, and stimulated with 1 μg/ml anisomycin for 60 min. The JNK KTR-mCherry and mKO-MK2 C/N ratios of single cells were plotted against basal nuclear mCherry and mKO intensities, respectively, on a logarithmic scale. The scatterplots were fitted to a linear regression and the obtained Pearson correlation values are shown. n ≥ 40 cells for each condition.

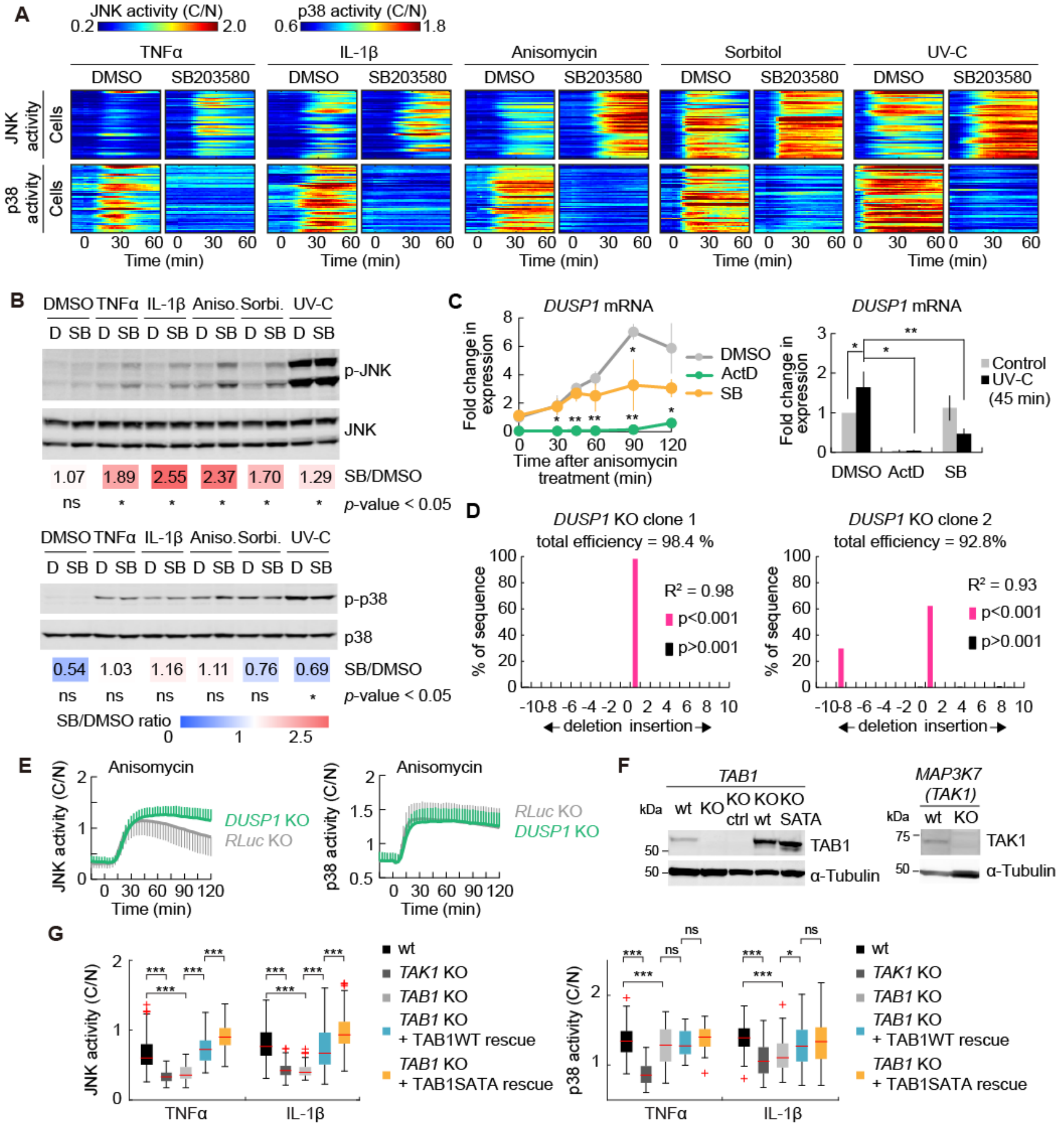


**Fig. S2. Cross-inhibition of JNK by p38.** (**A**) Clonal HeLa cells stably expressing JNK KTR-mCherry, mKO-MK2, and NLS-iRFP-NLS were pretreated with 0.1% DMSO and 10 µM SB203580, respectively, and then stimulated with 10 ng/ml TNFα, 10 ng/ml IL-1β, 10 ng/ml anisomycin, 200 mM sorbitol, or 100 J/m2 UV-C. JNK KTR-mCherry and mKO-MK2 C/N ratios are displayed as heat maps. Each line of the JNK KTR-mCherry and mKO-MK2 C/N ratios represents the activity dynamics in a single cell, with red colors indicating high and blue colors indicating low kinase activities. n = 50 cells for each condition from two independent experiments. (**B**) HeLa cells were pretreated with 0.1% DMSO (D) or 10 μM SB203580 (SB) and then stimulated with 10 ng/ml TNFα for 30 min, 10 ng/ml IL-1β for 60 min, 10 ng/ml anisomycin for 60 min, 200 mM sorbitol for 60 min, or 100 J/m2 UV-C for 60 min. Cell lysates were analyzed by western blotting against p-p38, p38, p-JNK, and JNK. Representative blots out of three independent experiments are shown. The SB/DMSO ratio is calculated as the fold change of the p-p38/p38 or p-JNK/JNK in the SB203580 over DMSO control treated samples. n=3 for each condition. Significance was tested by a Student's *t*-test. ns = not significant, *p<0.05. (**C**) HeLa cells were starved in imaging medium for 3 h and pretreated with 0.1% DMSO, 1 μg/ml actinomycin D, or 10 μM SB203580. mRNA levels of DUSP1 were measured in HeLa cells by qPCR after stimulation with 10 ng/ml anisomycin. (**D**) Validation of DUSP1 KO. Insertions and deletions at the DUSP1 target locus in clonal DUSP1 KO cells were identified using Sanger sequencing and the TIDE web tool. (**E**) HeLa DUSP1 KO and RLuc KO control cells stably expressing NJP were stimulated with 10 ng/ml anisomycin, and imaged over time. The time courses of the averaged JNK KTR-mCherry and mKO-MK2 C/N ratios with SD are shown. n = 50 cells for each condition from two independent experiments. (**F**) Western blot analysis of the expression levels of TAB1 in wildtype (wt) cells, clonal TAB1 KO cells, TAB1 KO cells stably expressing an empty control construct, TAB1 KO cells rescued by stable expression of TAB1-wt, and TAB1 KO cells stably expressing TAB1 S423A T431A (SATA). α-tubulin was used as a loading control (left). Western blot analysis of the expression levels of TAK1 in the wildtype (wt) and MAP3K7 KO cells (right). α-tubulin was used as a loading control. (**G**) HeLa wildtype, TAK1 KO, TAB1 KO cells, and TAB1 KO cells either complemented with TAB1 wildtype or TAB1 S423A T431A (SATA), stably expressing NJP were stimulated with 10 ng/ml TNFα for 30 min, or 10 ng/ml IL-1β for 50 min. JNK KTR C/N and mKO-MK2 C/N ratios are shown. The central line, top and bottom edges indicate the median, 25th, and 75th percentiles, respectively, with the whiskers denoting 1.5 times the interquartile range. Red crosses are outliers. n ≥ 34 cells for each condition from at least two independent experiments. Significance was tested against the wildtype control by a Student's *t*-test. ns, not significant; *p<0.05, **p<0.01, ***p<0.001.

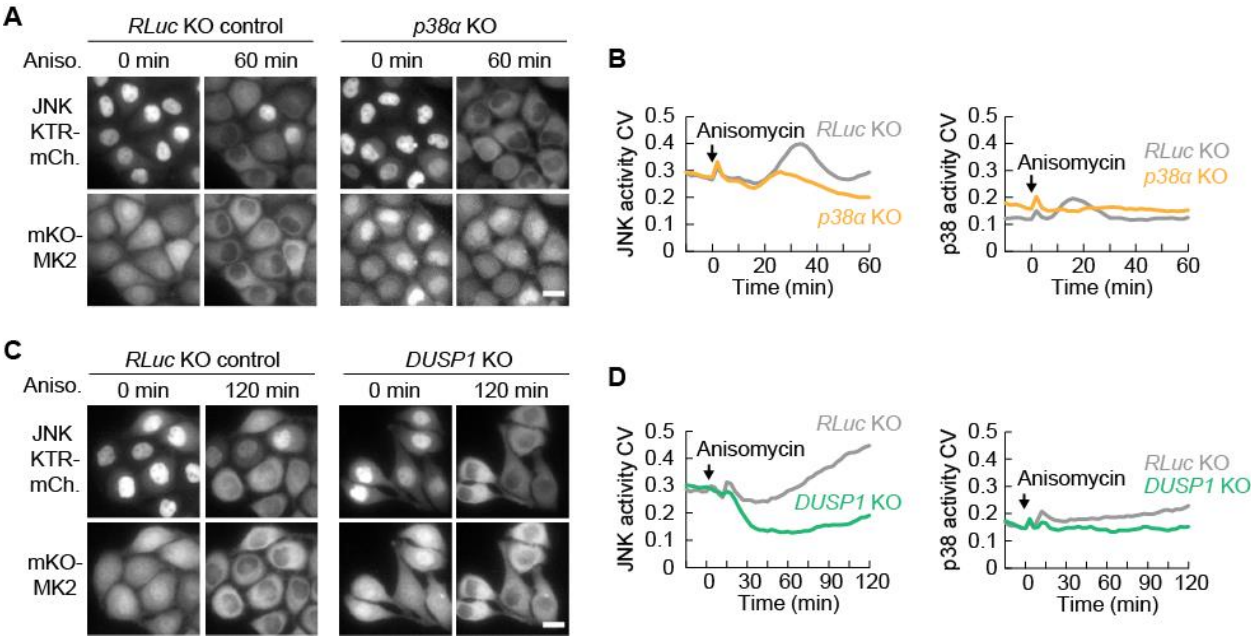
A
RLuc KO control
p38α KO
Aniso.
0 min
60 min
0 min
60 min
JNK KTR-mCh.
mKO-MK2
B
JNK activity CV
Anisomycin
RLuc KO
p38α KO
Time (min)
p38 activity CV
Anisomycin
RLuc KO
p38α KO
Time (min)
C
RLuc KO control
DUSP1 KO
Aniso.
0 min
120 min
0 min
120 min
JNK KTR-mCh.
mKO-MK2
D
JNK activity CV
Anisomycin
RLuc KO
DUSP1 KO
Time (min)
p38 activity CV
Anisomycin
RLuc KO
DUSP1 KO
Time (min)

**Fig. S3. Cell-to-cell variability of JNK and p38 activity upon inflammatory and stress stimuli.** (**A** and **B**) HeLa p38α KO and RLuc KO control cells stably expressing NJP were stimulated with 10 ng/ml anisomycin and imaged over time. Representative images of JNK KTR-mCherry and mKO-MK2 are shown (A). Scale bar: 20 µm. The time courses of the coefficients of variation (CV) of JNK KTR-mCherry and mKO-MK2 C/N ratios are represented. n = 110 cells for p38α KO and 120 cells for RLuc KO from two independent experiments. (**C** and **D**) HeLa *DUSP1* KO and *RLuc* KO control cells stably expressing NJP were stimulated with 10 ng/ml anisomycin and imaged over time. Representative images of JNK KTR-mCherry and mKO-MK2 before and 120 min after stimulation are shown (C). Scale bar: 20 µm. The time course of the CV upon 10 ng/ml anisomycin stimulation is shown for the JNK KTR-mCherry and mKO-MK2 C/N ratios in DUSP1 KO and RLuc KO control cells. n = 50 cells per condition from two independent experiments.

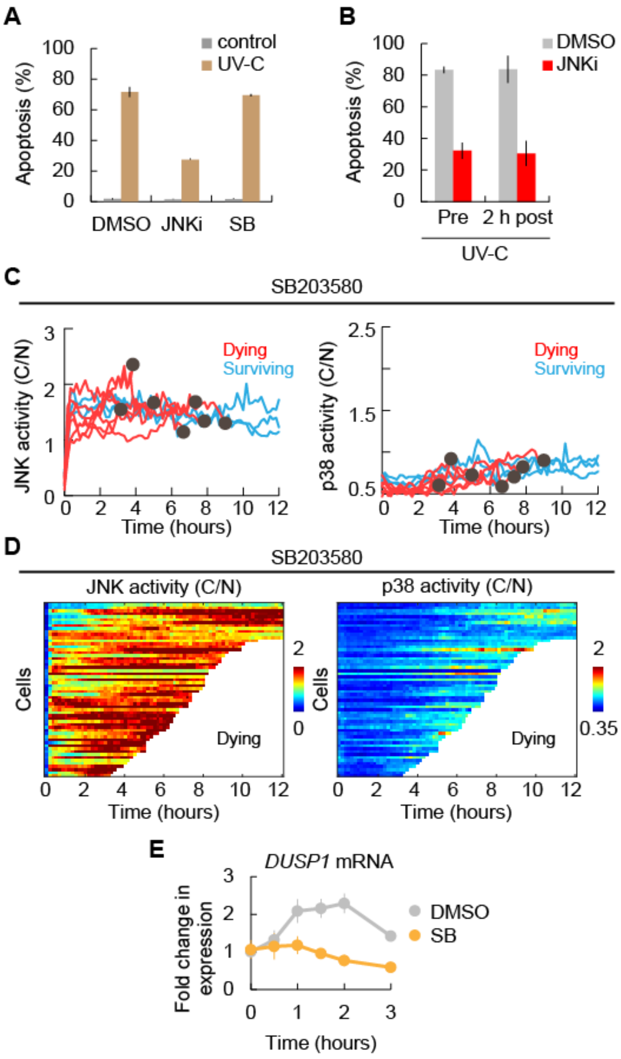


**Fig. S4. Cell-to-cell variation in JNK activity leads to fractional killing upon UV-C treatment.** (**A**) Apoptosis of 0.1% DMSO, 10 µM SB203580, or 10 µM JNK inhibitor VIII pretreated HeLa cells 12 h after 100 J/m$^2$ UV-C or mock treatment. Apoptosis was assayed by caspase 3 activation. Data are shown as the mean with SD. n = 3 independent experiments. (**B**) Apoptosis of 0.1% DMSO or 10 µM JNK inhibitor VIII treated HeLa cells 12 h after 100 J/m$^2$ UV-C. Cells were treated with inhibitor either before (pre) or 2 h after UV-C. Apoptosis was assayed by caspase 3 activation. Data are shown as the mean with SD. n = 3 independent experiments. (**C**) C/N ratios of JNK KTR-mCherry and mKO-MK2 in the presence of SB203580 were quantified for dying (red lines) and surviving (blue lines) cells after 100 J/m$^2$ UV-C stimulation. n = 10 representative cells. Black dots indicate apoptosis. (**D**) JNK KTR-mCherry and mKO-MK2 C/N ratios in the presence of SB203580 are displayed as heat maps for cells from (B). n = 58 cells. White traces indicate cells that underwent cell death. (**E**) Average fold changes of *DUSP1* mRNA levels of DMSO or 10 μM SB203580 pretreated HeLa cells in response to 100 J/$m^2$ UV-C, as measured by qPCR. n = 3 independent experiments.

**Movie S1. Multiplexed imaging of JNK and p38 activities upon anisomycin treatment.** HeLa cells stably expressing JNK KTR-mCherry (left), mKO-MK2 (middle), and NLS-iRFP-NLS (right) were imaged every 2 min. 10 ng/ml anisomycin was added at elapsed time point 0 min. Scale bar: 50 μm.

**Movie S2. Multiplexed imaging of JNK and p38 activities upon UV-C stress.** HeLa cells stably expressing JNK KTR-mCherry (left top), mKO-MK2 (right top), and NLS-iRFP-NLS (left bottom) were imaged every 10 min for 12 h following 100 $J/m^2$ UV-C exposure. Scale bar: 50 μm.
